# Estimating the impact of invasive pests and diseases on ecosystem services: modelling carbon sequestration loss due to myrtle rust (*Austropuccinia psidii* exotic strains) in Australia

**DOI:** 10.1101/2025.05.30.657121

**Authors:** Thao P. Le, Meryl Theng, Chris M. Baker, Isobel R. Abell, Tom Kompas, Emma J. Hudgins

**Affiliations:** The Centre of Excellence for Biosecurity Risk Analysis, The University of Melbourne, Grattan St, Parkville, 3010, Victoria, Australia; School of Mathematics and Statistics, The University of Melbourne, Grattan St, Parkville, 3010, Victoria, Australia; Melbourne Centre for Data Science, The University of Melbourne, Grattan St, Parkville, 3010, Victoria, Australia; School of Agriculture, Food and Ecosystem Sciences, The University of Melbourne, Grattan St, Parkville, 3010, Victoria, Australia

**Author notes:** These authors contributed equally to this work (Thao P. Le, Meryl Theng). Email addresses:* (Thao P. Le), (Meryl Theng).

**Keywords:** carbon accounting, ecosystem services, environmental valuation, forest pathogen, ecological and economic impacts

## Abstract

The impacts of invasive pests and diseases are routinely estimated and measured in the context of agriculture, but less so in the context of biodiversity and ecosystem services. In this study, we estimate the potential reduction of carbon sequestration in Australia due to exotic strains of myrtle rust (*Austropuccinia psidii*, also known as *Puccinia psidii*, guava rust, or ōhi’a rust). We model the contribution of susceptible plants to carbon sequestration and use previously known myrtle rust damage estimates to susceptible plant species and the valuation of carbon sequestration in Australia to estimate the potential monetary impact. This method can be systematically extended to other pests impacting plant growth as well as other ecosystem services. In the case of myrtle rust, we estimate that it could cause up to a 1.6% (95% CI: 1.3–2.0%) annual reduction in national carbon sequestration if it were to spread across all climatically suitable areas in Australia, resulting in an estimated value loss of over $340 million AUD (over $220 million USD) per year. Compared with contemporary syntheses of known cost estimates, our results show that the potential consequences of invasive species can be substantially larger than reported, and may be currently undervalued. Our work shows the need to systematically compile the potential impacts and costs to the environment and ecosystem services globally, to support both biosecurity decision-making and climate-change related initiatives such as net-zero emissions targets and reforestation efforts.

## 1. Introduction

Biodiversity provides crucial ecosystem services, including clean air, food, water, defence against natural disasters, as well as cultural benefits to humans [1, 2, 3]. Given the use of economic costs as a common currency to assess a range of negative consequences to human and natural systems, ascribing value to the environment — and the potential devastating impacts from human-mediated threats — provides an explicit rationale for environmental protection through the enumeration of ecosystem services [4, 5, 6, 7, 8, 9]. Environmental valuation and impact estimation enforces transparency in decision-making around threats to these services and quantifies the risk and consequences of failing to protect these services. Given the available literature on ecosystem valuation [10] (including for Australia for multiple ecosystem services [7]), we will be focused on the impacts and losses due to threats, in particular due to *invasive pests and diseases*.

Human movement and global trade has accelerated the spread of invasive pests and diseases due to the facilitation of long-distance travel [11]. According to the Intergovernmental Platform on Biodiversity and Ecosystem Services [12], invasive alien species are one of the top five threats to biodiversity (alongside changes in land and sea use, direct exploitation of organisms, climate change, and pollution). There are numerous studies investigating the potential impact of invasive species on non-provisioning ecosystem services and the environment (for example, [13, 14, 15] and citations within), but far fewer including cost estimates (e.g., [8, 9, 16]). The most recent compilation of invasive species economic impacts is the database ‘InvaCost’ [8], which is a global register of cost estimates and reports for all invasive species globally. However, this database focuses primarily on agricultural costs — reflecting the greater cost-estimation work done on invasive species impacts to industry and the agricultural sector [17, 18] — and has fewer recorded observed or potential costs for non-market impacts, such as impacts to the environment; it records a $600 billion USD impact since 1970 to agriculture, fisheries and forestry [6], while estimating only $50 billion USD impact to the environment globally. For comparison, in the case of Australia, Stoeckl at al [7] estimated a smaller disparity between the potential impacts on portfolio assets (including agriculture, fisheries and forestry) and regulating ecosystem services (and water) due to unmitigated spread of invasive invertebrates and weeds ($24.4 billion versus $7.9 billion AUD respectively).

Note that this is still orders of magnitude smaller than the estimated loss of $4.3–20.2 trillion per year due to land-use changes between 1997 to 2011 [19]. In sum, we suspect that the environmental costs of invasive species, in InvaCost and in general, may be underestimated due to lack of systematic compilation. Given the increasing spread and risk of invasive pests and diseases, which can be exacerbated by land-use changes and climate change [20], a systematic compilation of potential impacts due to invasive species on ecosystem services is needed to support environmental protection via biosecurity measures [21].

There are three broad methods for estimating the potential impacts of new invasive species to ecosystem services and the environment. Firstly, there are survey methods such as structured expert elicitation [14, 22, 23] and choice modelling [16]. Secondly, there are benefit transfer methods, which take previous studies of impacts in other locations and estimate impact at a new site [7, 14, 23], taking into account factors such as GDP and population density. Thirdly, there are modelling methods that consider the mechanisms of impact and cost [9, 24]. Prior systematic compilation of potential impacts to the Australian environment in particular employed expert elicitation and benefit transfer [14]. While expert elicitation methods are particularly useful when qualitative information is limited, they cannot directly harness the increasing scientific data that *are* available. However, given the complexity and individuality inherent in mechanistic modelling, it may not be feasible to use mechanistic modelling to systematically estimate the potential impacts of dozens, if not thousands, of invasive species [25] across multiple ecosystem services. Refs. [7, 23] take a mixed approach, using expert elicitation to estimate the potential impact of invasive species (at a functional group level) on ecosystem services, and then modelling the spread of those invasive species across Australia into vulnerable regions to estimate the overall potential impact on ecosystem services.

Here, we develop a parsimonious modelling workflow to estimate potential environmental impacts due to invasive pests and diseases, which we term “contribution modelling”. The approach is applied here in a case study, but could be applied systematically across invasive species and ecosystem services to estimate their total potential impacts at the country scale. Contribution modelling involves assessing the makeup of susceptible and non-susceptible species to an ecosystem service, incorporating host-specific invasive pest/disease impacts to each host species’ capacity to contribute to that ecosystem service, and combining that with the economic evaluations of the ecosystem service to estimate potential impacts at the country scale.

As a case study, we estimated the impact of exotic strains of myrtle rust (*Austropuccinia psidii*) on carbon sequestration in Australia. Myrtle rust is a fungal disease that affects eucalypts, a common tribe of Australian tree genera that perform substantial amounts of Australia’s forest carbon sequestration and storage. According to Australian *National Priority List of Exotic Environmental Pests, Weeds and Diseases* [14], myrtle rust is expected to have a ‘massive’ impact on the environment. Given the current spread of the pandemic (i.e., established) strain of myrtle rust in Australia and the large amount of information regarding other species’ susceptibility to myrtle rust [26], the impact of not-yet-established strains of myrtle rust (exotic strains) was deemed an ideal and high-priority case study.

We chose to focus on estimating ecosystem service losses from carbon sequestration. This service is arguably easier to estimate than most other ecosystem services, given the existence of the carbon market and the more direct relationship between invasive pest and disease impact and subsequent reduction in carbon sequestration. There are a wide spectrum of plant diseases and plant pests that can stunt plants and trees and thus reduce carbon sequestration; in the extreme cases, they can cause tree death, corresponding to a complete loss in carbon sequestration ability from that tree. Work evaluating past impact of invasive pests and diseases on carbon sequestration exist (e.g., [27, 28, 29, 30, 31, 32, 33] and references therein), but much fewer works estimate potential future impacts from exotic pests and diseases [24, 34, 35] (noting that these do not include economic dollar estimates). Of the entries filed under having terrestrial environmental impacts in the InvaCost database, only 19 entries (out of a total 13554 in version 4.1) mention ecosystem services or carbon sequestration [8].^1^ In contrast, there have been numerous global studies studying the future impacts on carbon sequestration due to threats such as climate change and human-mediated deforestation and landuse changes (e.g. [36, 37, 38, 39, 40, 41, 42, 43, 44] and references therein). Given that carbon sequestration is fundamental to climate-change mitigation efforts, but there is a lack of systematic estimation of the potential impacts from invasive pest and diseases, it was deemed a fitting ecosystem service to focus on for the case study.

To estimate the potential impact of myrtle rust (exotic strains) on Australian carbon sequestration, we begin by employing national ecological datasets to estimate the distribution of plants, at the genus level, across Australia’s Major Vegetation Groups (MVG) [45]. Using available biometric information and average growth, we calculate genus-level contributions to carbon capture across Australia, assuming above-ground and below-ground biomass within terrestrial vegetation classes. We also use a range of scientific studies to determine the impacts of myrtle rust at the species level, and bioclimatic variables to delineate areas with myrtle rust infection risk. By combining the estimated loss due to myrtle rust impacts with previously estimated valuations carbon sequestration in Australia [7], and assuming a linear relationship between amount of carbon sequestered and value, we are able to estimate the total loss in terms of Australian dollars.

## 2. Materials and Methods

### 2.1. Modelling framework overview

There are three key steps to our modelling framework to estimate the impact of invasive pests and diseases on ecosystem services:

1. Estimate the current spatial distribution of ecosystem service provision, which involves substeps:
  a. Estimating the spatial distribution of the ecosystem service provider(s)
  b. Estimating the current ecosystem service contribution of each provider
2. Estimate the potential impact of invasive species on each provider’s provision of the ecosystem service, by:
  a. Estimating the spatial risk distribution of the invasive species
  b. Estimating the individual impact of the invasive species on each provider type’s ability to contribute to the ecosystem service.
3. Estimate the potential monetary damages on the ecosystem service by the invasive species, by:
  a. Estimating the current valuation of the ecosystem service
  b. Estimating the potential damage by assuming a relationship between provision and valuation.

Here, we consider the impact of myrtle rust (exotic strains) on carbon sequestration. As such, the steps are:

1. Estimate the current spatial distribution of carbon sequestration by woody plants in Australia, by:
  a. Estimating the spatial distribution of woody plants, and
  b. Estimating the annual carbon sequestered by woody plants within each genus.
2. Estimate the potential impact of myrtle rust (exotic species) on carbon sequestration, by:
  a. Estimating the spatial climatic infection risk of myrtle rust in Australia, and
  b. Estimating the impact of myrtle rust on the ability of woody plant species within each genus to sequester carbon.
3. Estimate the potential monetary damages on the ecosystem service by the invasive species, by:
  a. Estimating the current valuation of carbon sequestration (which has already been estimated by [7]), and
  b. Estimating the potential damage by assuming a linear relationship between annual carbon sequestration and annual carbon sequestration valuation.

A flow chart of the different components of the modelling process for the case study is presented in Figure 1. Note that modelling spread is not explicitly part of this modelling framework, but the results of contribution modelling can be embedded into existing models of invasive species spread [7, 23, 46].

**Figure 1.**
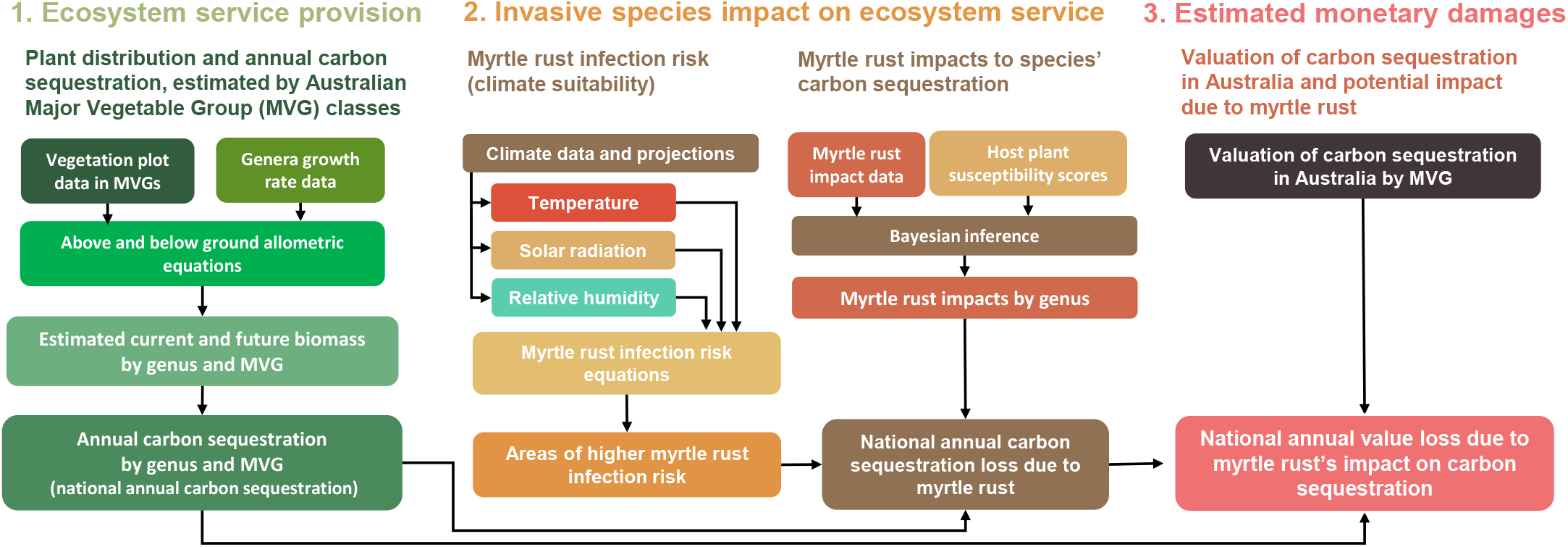
Flow chart of the process for estimating the potential monetary impact of myrtle rust (exotic strains) on carbon sequestration in Australia.

### 2.2. Study area

Our approach estimates country-scale impacts on ecosystem services, which we illustrate with our Australian case study. With a total area of 7.7 million km^2^ [47], Australia has 6 climatic zones based on temperature and humidity [48], 32 Major Vegetation Groups (MVG) based on dominant cover and vegetation distribution [49], and 89 geographically distinct bioregions based on climate, geology, landform, and vegetation and species distribution [50]. According to the Australian Bureau of Agricultural and Resource Economics and Sciences (ABARES), Australia has approximately 133.6 million hectares of forest (17% of Australia’s land area, as of 2021) [51], with Eucalypt forest comprising 77% of its total native forest area (101.1 million hectares) [51].

For modelling purposes, in this study we subdivide Australia by the major vegetation groups [49]: (1) Rainforests and vine thickets, (2) Eucalypt tall open forests, (3) Eucalypt open forests, (4) Eucalypt low open forests, (5) Eucalypt woodlands, (6) Acacia forests and woodlands, (7) Callitris forests and woodlands, (8) Casuarina forests and woodlands, (9) Melaleuca forests and woodlands, (10) Other forests and woodlands, (11) Eucalypt open woodlands, (12) Tropical eucalypt woodlands/grasslands, (13) Acacia open woodlands, (14) Mallee woodlands and shrublands, (15) Low closed forests and tall closed shrublands, (16) Acacia shrublands, (17) Other shrublands, (18) Heathlands, (19) Tussock grasslands, (20) Hummock grasslands, (21) Other grasslands, herblands, sedgelands and rushlands, (22) Chenopod shrublands, samphire shrublands and forblands, (23) Mangroves, (24-30) Other cover types, (31) Other open woodlands, and (32) – Mallee open woodlands and sparse mallee shrublands.

### 2.3. Exemplar invasive species

In this study, we consider the impact of myrtle rust. Myrtle rust is disease of plants caused by the fungus *Austropuccinia psidii*. It can cause deformed leaves, branch defoliation, dieback, stunted growth, and even plant death [26, 52]. It affects plants belonging to *Myrtaceae* family, including eucalypts, willow myrtles, bottlebrushes and paperbarks, with over 200 species confirmed infected [26] and over 300 determined susceptible [53] in Australia. It can spread via wind, rain, animals and by human movement [54]. Since its first detection in Australia in 2010 [55], the ‘pandemic’ strain of myrtle rust has since spread across New South Wales, Victoria, Queensland, Tasmania, the Tiwi Islands in the Northern Territory and the northern part of Western Australia [56]. Several other strains of myrtle rust exist with varying virulence and species impacts [57]. Here, we model the potential impact of the invasion and countrywide spread of a hypothetical exotic strain of myrtle rust that is equally impactful, if not worse, than the currently established strain.

### 2.4. National spatial carbon contribution by plant genus

Carbon sequestration is the rate at which an organism or ecosystem is able to store carbon, often measured annually. Our approach starts by estimating the amount of dry woody biomass across Australia, and combines this information with plant growth models to estimate countrywide annual carbon sequestration. Estimating carbon sequestration or storage is rarely done with any amount of taxonomic granularity, and is most often reported at an ecosystem level (e.g., [58, 59, 60, 61]), in part due to the high data requirements (i.e., plot-level vegetation distributions) of finer-scale estimates. In our case, we overcome this burden by making the assumption that vegetation within a major vegetation group (MVG) is homogeneous. We use detailed survey plots containing plant composition and diameter information for each MVG to determine average plant distributions and estimate annual tree growth using empirical estimates available at the genus level, allowing us to estimate annual carbon sequestration for each MVG. The steps are:

1. Estimate the biomass per tree within the vegetation survey via its ‘current’ diameter measurement,
2. Predict the growth in diameter of each tree in one year’s time,
3. Estimate the biomass per tree in a year’s time based on the ‘future’ diameter (hence accounting for growth),
4. Subtract the ‘current’ biomass from the ‘future’ biomass to obtain an estimate of biomass sequestered per tree per year.
5. Repeat this for every tree in the survey plot, and extrapolate biomass across the major vegetation group area.
6. If multiple surveys exist in the same MVG, we average across the surveys.
7. Combine information across all MVGs to obtain national estimates of carbon sequestration.

This process allows us to obtain MVG-specific percentage contributions of each major plant taxon. We estimate contribution at the genus level, as species-level information within MVG survey plots was not always available. Genus (or species)-level carbon sequestration is a key innovation of our workflow, as it allows us to directly estimate the potential spatial impacts from myrtle rust on specific genera.

#### 2.4.1. Spatial distribution of trees and shrubs

We collated openly available Australian vegetation plot datasets of individual trees and shrubs that contained species and diameter information of each plant record, and coordinates of the plot or site, sourcing them from the *Biomass Plot Library* [45], *Calperum Mallee Stem Diameter, Height and Aboveground Woody Biomass Data* [62] and *Robson Creek Rainforest Diameter, Height and Aboveground Woody Biomass Data* [63].

Note that we discarded records before 2013 and assumed that the dataset was representative of the present vegetation within the surveyed plots. In addition, we discarded records without species or diameter information, and only kept the latest record for individuals that were recorded more than once.

We then extrapolated plots within their MVG and assumed homogenous MVGs. For MVGs with no data, we were unable to estimate their carbon sequestration by biomass; as a conservative estimate, we assumed that those MVGs had no biomass loss, hence no carbon sequestration value loss.

#### 2.4.2. Biomass estimation

We use allometric equations based on plant functional types to estimate an individual tree or shrub’s biomass using diameter, height, species, region, and other tree and stand characteristics. There are separate allometric equations for above-ground biomass (AGB) and below-ground biomass (BGB) [64, 65]. To estimate above-ground biomass, individual tree and shrub records were first grouped into plant functional types shrubs, multistemmed hardwood trees, single-stemmed hardwood trees, and other trees, before applying the relevant allometric equations from Paul et al [64] to predict aboveground biomass. Similarly, to estimate below-ground biomass, individual tree and shrub records were grouped into the plant functional types: shrubs, multi-stemmed trees, single-stemmed trees and trees of relatively low stem woody density, before applying the relevant allometric equations from Paul et al [65] to predict below-ground biomass. The total biomass from a particular tree or shrub is the combination above- and below-ground biomass. Further details are provided in Appendix A.2.

#### 2.4.3. Carbon sequestration per year

To estimate carbon sequestration rate, we first estimated the growth in biomass for each individual tree or shrub *i*. We compiled species- or genus-specific annual diameter growth rates from several sources [66, 67, 68, 69]. We then added the annual growth rate to the existing diameter measurement for the plant record to obtain a “future” diameter measurement (i.e., growth in diameter after one year). The “future” biomass was estimated from the “future” diameter estimates using the AGB and BGB allometric equations. We then calculated the biomass growth (after one year) for each plant record *i* by subtracting the “current” biomass at time *t* (*B*_*i,t*_) from the “future” biomass estimate at time *t* + 1 (*B*_*i,t*+1_). Finally, carbon sequestration (*C*_*i*_) rate per plant was obtained by multiplying biomass growth by 50%:

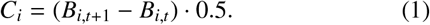

This was based on the widely accepted assumption that 50% of dry woody biomass is carbon [70, 71].

Note that we assumed that trees from the same genus grow at the same annual genus mean growth rate (i.e., averaged across species, ages, climate). Genera lacking diameter growth rate information (e.g., *Acmena, Astrobuxus, Archirhodomyrtus, Nothofagus, Nematolepis, Olearia*) were conservatively assumed to have a slow growth rate of 0.1 cm/year. Trees or shrubs with *D*_10_ measurements (i.e., diameter measured at 10 cm from the base) were discarded from the analysis due to the lack of growth information at that height measurement. These trees formed ∼ 12% of total biomass in our dataset.

#### 2.4.4. Carbon contribution per major vegetation group per hectare

For each major vegetation group, the carbon contribution of each genus *g* (i.e., the amount of annual sequestration by each genus) was estimated by taking the sum of carbon sequestered by plant records *i* of the same genus (∑ _*i* if genus *g*_ *C*_*i*_) and dividing it by the sum of all areas *A*_*x*_ (in hectares) of sampled plots *x* in the relevant MVG (∑ _*x* in MVG_ *A*_*x*_). This derives a per hectare per year (tonnes ha^™1^ y^™1^) estimate of carbon sequestration for each genus:

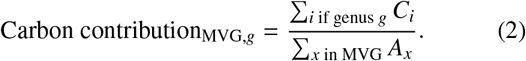

### 2.5. Potential impact of myrtle rust on carbon sequestration ability

To estimate the potential impact of myrtle rust on carbon sequestration of susceptible species, we first modelled regions in Australia at risk of infection, using physiological parameters and bioclimatic variables. Spatial infection risk was assessed under three climate models, each estimating current conditions and projecting three future climate scenarios to capture uncertainty. Estimates of impacts of myrtle rust to different species from the literature were aggregated to the genus level. Genus-specific impact values were then applied to adjust (i.e., reduce) the carbon sequestration contribution of susceptible trees and shrubs within at-risk areas.

#### 2.5.1.Myrtle rust infection risk

We estimated regions at risk of myrtle rust infection using the infection risk formula of Beresford et al [72]. Myrtle rust infection risk, *I*_risk_, is predicted using bioclimatic variables including temperature (mean temperature during the period of high-relative-humidity (RH), i.e., relative humidity ≥ 85%), the number of hours with high relative humidity, and hourly solar radiation.

Note that we rely on quarterly temperature data from the three-month period with the highest precipitation in each raster pixel. Given our lack of hourly temperature data, we assume that daily temperatures have a cosine form (Eq. (7) in [73]) and assume that maximum temperatures occur at 3 pm and minimum temperatures occur at 6 am [74]. Furthermore, to estimate hours of high relative humidity in each day, we first estimated the hourly relative humidity based on hourly and daily saturation vapour pressure, temperature, and daily humidity. See Appendix A.3 for further details.

After calculating the myrtle rust infection risk *I*_risk_ (Eq. (A.19)) across Australia, we classified suitable areas using two thresholds: *I*_risk_ > 0.2 (less conservative, assuming greater spread) and *I*_risk_ > 0.5 (more conservative, assuming less spread). Higher thresholds were not chosen as they were too conservative and do not reflect the current known distribution of myrtle rust (pandemic strain) in Australia. These masks were subsequently used to estimate the potential impact of myrtle rust on vegetation.

The bioclimatic variables required to compute *I*_risk_ were derived from CMIP6 (Coupled Model Intercomparison Project) climate model data, specifically: surface air temperature (variable `tas`), minimum temperature (`tasmin`), maximum temperature (`tasmax`), precipitation (`pr`), surface solar radiation (`rsds`), and relative humidity (`hurs`).

##### 2.5.2. Risk under various climate projections

To account for the uncertainty of the climate suitability at the present date, and to examine possible scenarios of future myrtle rust infection, we considered bioclimatic data from a range of CMIP6 climate models: ACCESS-CM2 (Australian Community Climate and Earth System Simulator, version 2, CCAM coupled ocean version), CNRM-CM6-1-HR (Centre National de Recherches Météorologiques Coupled Global Climate Model, version 6.1, high resolution), and MPI-ESM1-2-LR (Max Planck Institute Earth System Model, version 1.2, low resolution). These models were selected based on skill scores for Australia (i.e., ACCESS-CM2; [75]), the ability to obtain high-resolution (i.e., ACCESS-CM2, CNRM-CM6-1-HR), diverse predictions, and their inclusion of projections of the bioclimatic variables required to calculate myrtle rust infection risk. We considered these models at the ‘present’ time point (year 2014) and into the future (year 2100) under Shared Socioeconomic Pathways climate scenarios SSP1-2.6 (low emissions), SSP2-4.5 (moderate emissions) and SSP3-7.0 (high emissions) [76]. All data were sourced as monthly records from Ref. [77].

##### 2.5.3. Effects of myrtle rust on plant carbon sequestration

Myrtle rust affects tree growth, and can cause branch death and tree death, thus reducing or nullifying the carbon sequestration capacity of an infected tree. In order to determine the carbon sequestration reduction due to myrtle rust, we consider the following myrtle rust impacts: tree death, percentage branch death, and reduced tree growth (height) compared to uninfected trees.

Tree death corresponds to a 100% impact on carbon sequestration. If, for example, 50% of trees of a particular species die due to myrtle rust, we assume that myrtle rust has a 50% impact on carbon sequestration of that species. We assume that the physical action of carbon sequestration occurs through branches and that branches contribute equally to the sequestration of a tree. Thus, if a tree has 20% branch death, then there is a 20% impact on carbon sequestration. Finally, a tree sequesters carbon as it grows, and thus a reduced growth rate means it is sequestering carbon slower. If an infected tree is 10% shorter than its non-infected counterpart, then we assume that there is a 10% impact on carbon sequestration, as a simplification.

Table A.3 in the appendix lists the species with some known quantitative information regarding myrtle rust impacts, and their source. Ideally, we would have quantitative data on myrtle rust impacts for all plant genera across all MVGs, but these data are incomplete. However, there are compilations of qualitative susceptibility ratings [26, 78, 79]. We used the susceptibility ratings from Makinson [26] (Appendix 3 of [26]), along with the quantitative data (Table A.3), to estimate the percentage damage impact given some assigned rating using Bayesian inference.

The ratings we use are *relatively tolerant, moderate susceptibility, high susceptibility*, and *extreme susceptibility*. Figure A.14 shows a plot of the distribution of ratings from Makinson [26]. Given that non-linear nature of these categories, we use Bayesian inference to estimate the boundary of these ratings, i.e., the percentage damage range that each rating corresponds to, as well as the distribution of potential impacts of species across the damage spectrum.

Figure 2 shows an example of a set of parameters drawn from the posterior. For example, a species with rating “moderate susceptibility” will be estimated to have a loss of carbon sequestration loss between [θ_*RT*_, θ_*MS*_]. Damage parameters are drawn randomly per the distribution for subsequent calculation. See Appendix A.5 for the details.

**Figure 2.**
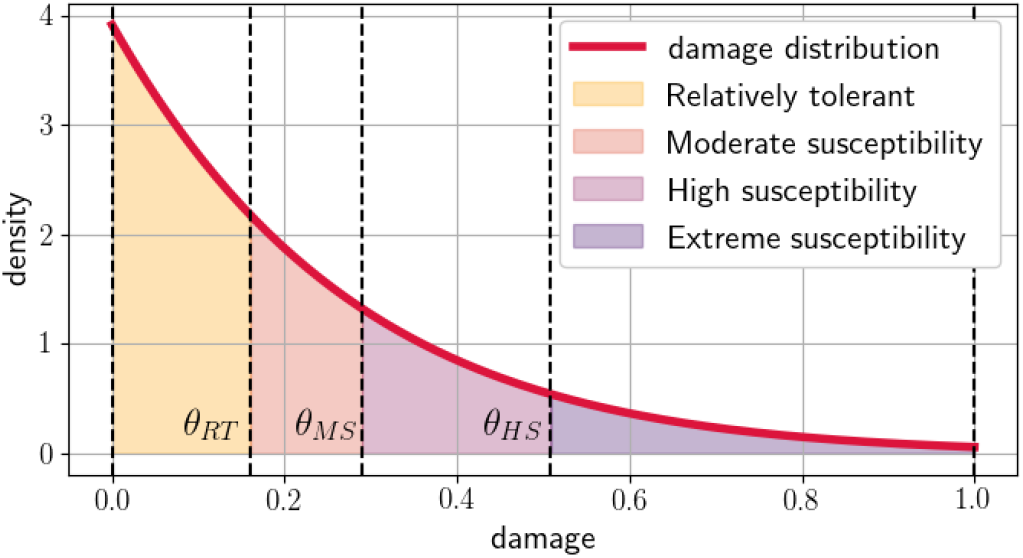
Estimating a damage value given a susceptibility rating: Example distribution of probability density of damage and thresholds drawn from the posterior distribution after Bayesian inference. The thresholds θ_*RT*_, θ_*MS*_, θ_*HS*_, and 1.0 mark the upper boundaries different susceptibility categories, relatively tolerant, moderate susceptibility, high susceptibility, and extreme susceptibility, respectively.

Using this method, we can obtain estimated myrtle rust damages on plant genera *Agonis, Asteromyrtus, Austromyrtus, Backhousia, Callistemon, Chamelaucium, Corymbia, Decaspermum, Eucalyptus, Eugenia, Gossia, Homoranthus, Hypocalymma, Lenwebbia, Leptospermum, Lindsayomyrtus, Lithomyrtus, Lophostemon, Melaleuca, Metrosideros, Mitrantia, Myrciaria, Myrtus, Neofabricia, Osbornia, Pilidiostigma, Rhodamnia, Rhodomyrtus, Ristantia, Sphaerantia, Stockwellia, Syzygium, Thryptomene, Tristania, Tristaniopsis, Uromyrtus*, and *Xanthostemon*.

##### 2.5.4. Potential impact of myrtle rust on carbon sequestration

We estimated the potential carbon sequestration reduction due to myrtle rust by major vegetation group. Given the carbon sequestration by genus by MVG from Sec. 2.4, we selected the subset of areas within each MVG that had a myrtle rust infection risk greater than some threshold (*I*_risk_ > 0.2 or > 0.5) (Sec. 2.5.1). Within these areas at risk, we applied the genus-specific myrtle rust percentage impact values on the carbon sequestration contribution of each genera.

Aggregating across genera and MVG, we can obtain the landscape-level loss of carbon sequestration. The percentage carbon reduction in each MVG is a simple ratio of the loss and original carbon sequestration value.

### 2.6. Economic valuation of potential carbon sequestration loss due to myrtle rust

#### 2.6.1. Valuation of carbon sequestration

The Australia-wide valuation of carbon sequestration was estimated by Stoeckl et al. [7, 80]. They estimated the 2015-AUD dollar value of carbon sequestered per hectare, across different major vegetation groups and natural resource management (NRM) regions using benefit transfer functions. See [7] (including Table C.1 of [7]) for more details on their process of carbon sequestration valuation.

The total annual value of carbon sequestration across Australia is estimated to be $22.6 billion, with major MVG contributors being *Eucalypt Open Forests* ($3.7 billion), *Eucalypt Woodlands* ($3.5 billion), and *Hummock Grasslands* ($2.8 billion).

#### 2.6.2. Valuing the potential impact of myrtle rust on carbon sequestration

To obtain the overall dollar value loss (or damage) due to myrtle rust, we assumed a proportional relationship between the amount of carbon sequestered and the value of carbon sequestration by MVG. Hence, a 10% loss in carbon sequestration in an MVG was calculated to be a 10% loss in carbon sequestration value of that MVG. For MVGs with no biomass data, we conservatively assumed that they will have no loss in carbon sequestration value due to myrtle rust. The national potential dollar impact of myrtle rust on carbon sequestration is then calculated as a sum of value losses across MVGs.

## 3. Results

### 3.1. Australian carbon sequestration by genus by major vegetation group

The collation of Australian vegetation plot data produced an inventory of 28,355 unique trees across 159 survey plots, spanning 50.5 hectares and representing 13 of the 32 Major Vegetation Groups (MVGs; Table A.2). Based on this dataset, *Eucalyptus* species were estimated to be the dominant contributors to carbon sequestration, accounting for 50–100% of the carbon sequestered by woody plants in 12 of the 13 MVGs assessed. See Fig. 3 for the carbon sequestered by genus by MVG.

**Table A.2:** Summary of plot data collated for each Major Vegetation Group.

| Major Vegetation Group | No. of plots | Total area surveyed (ha) | No. of trees |
| --- | --- | --- | --- |
| 1 Rainforests and Vine Thickets | 2 | 2 | 1620 |
| 2 Eucalypt Tall Open Forests | 52 | 18.26 | 11148 |
| 3 Eucalypt Open Forests | 40 | 14.61 | 7867 |
| 4 Eucalypt Low Open Forests | 1 | 1 | 795 |
| 5 Eucalypt Woodlands | 18 | 4.7 | 1517 |
| 6 Acacia Forests and Woodlands | 0 | 0 | 0 |
| 7 Callitris Forests and Woodlands | 1 | 0.25 | 254 |
| 8 Casuarina Forests and Woodlands | 0 | 0 | 0 |
| 9 Melaleuca Forests and Woodlands | 0 | 0 | 0 |
| 10 Other Forests and Woodlands | 0 | 0 | 0 |
| 11 Eucalypt Open Woodlands | 9 | 0.32 | 86 |
| 12 Tropical Eucalypt Woodlands/Grasslands | 0 | 0 | 0 |
| 13 Acacia Open Woodlands | 0 | 0 | 0 |
| 14 Mallee Woodlands and Shrublands | 2 | 2 | 89 |
| 15 Low Closed Forests and Tall Closed Shrublands | 0 | 0 | 0 |
| 16 Acacia Shrublands | 1 | 0.06 | 91 |
| 17 Other Shrublands | 0 | 0 | 0 |
| 18 Heathlands | 0 | 0 | 0 |
| 19 Tussock Grasslands | 1 | 0.04 | 6 |
| 20 Hummock Grasslands | 0 | 0 | 0 |
| 21 Other Grasslands, Herblands, Sedgeland and Rushlands | 1 | 0.04 | 20 |
| 22 Chenopod Shrublands, Samphire Shrublands and Forblands | 1 | 0.04 | 1 |
| 23 Mangroves | 0 | 0 | 0 |
| 24 Inland Aquatic | 0 | 0 | 0 |
| 25 Cleared, non-native veg., buildings | 30 | 6.16 | 4861 |
| 26 Unclassified native veg. | 0 | 0 | 0 |
| 27 Naturally bare - sand, rock, claypan, mudflat | 0 | 0 | 0 |
| 28 Sea and estuaries | 0 | 0 | 0 |
| 29 Regrowth, modified native vegetation | 0 | 0 | 0 |
| 30 Unclassified forest | 0 | 0 | 0 |
| 31 Other Open Woodlands | 0 | 0 | 0 |
| 32 Mallee Open Woodlands and Sparse Mallee Shrublands | 0 | 0 | 0 |

**Figure 3.**
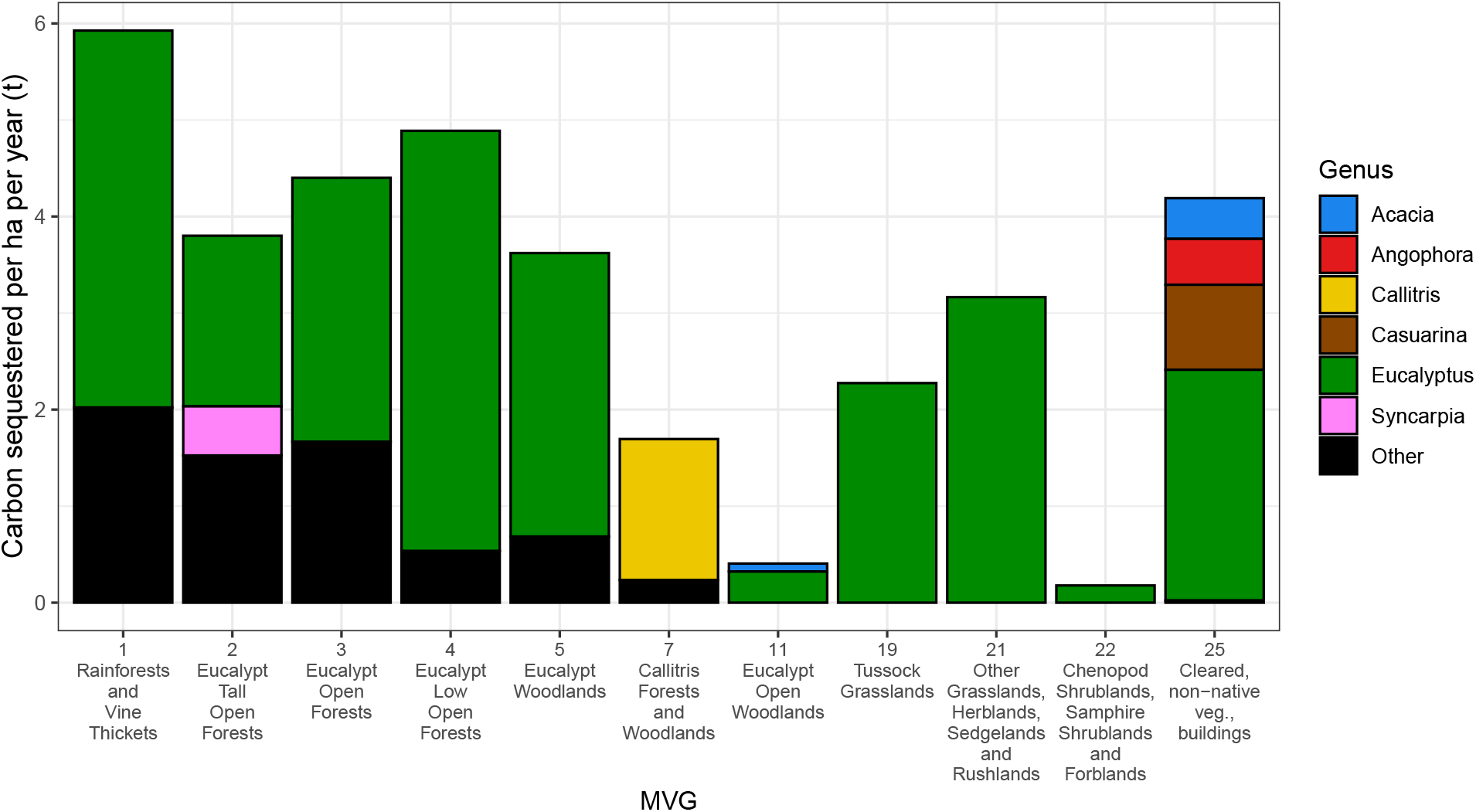
Carbon sequestered by each genus per hectare per year (tonnes ha^™1^ y^™1^) within each Major Vegetation Group (MVG). Only the top contributing plant genera are shown and all other genera (<10% contribution) are grouped under “Other”.

Note that of the MVGs assessed, eight were represented by two plots or fewer, covering no more than two hectares in total. In addition, no data were available for 19 of the 32 MVGs.

### 3.2. Myrtle rust impacts on genus

Table 1 shows the median and 95% credible interval on the estimated impacts of myrtle rust on different genera. All other genera not listed are assumed to be not susceptible to myrtle rust, as a conservative estimate. Top relevant impacts are to *Eucalyptus* (22% [18%, 27%]), *Syzygium* (13% [10%, 16%]) and *Rhodamnia* (37% [32%, 42%]).

**Table 1:** Estimated impacts of myrtle rust on different genera.

| Genus | Median | 95% credible interval |
| --- | --- | --- |
| *Acacia* | 0.0% | [0.0%, 0.0%] |
| *Agonis* | 64.6% | [50.4%, 94.6%] |
| *Archirhodomyrtus* | 94.3% | [94.3%, 94.3%] |
| *Asteromyrtus* | 4.8% | [0.2%, 13.0%] |
| *Austromyrtus* | 10.1% | [1.7%, 27.3%] |
| *Backhousia* | 12.6% | [8.1%, 18.1%] |
| *Baeckea* | 11.0% | [11.0%, 11.0%] |
| *Callistemon* | 19.9% | [14.2%, 26.3%] |
| *Chamelaucium* | 63.9% | [50.6%, 94.1%] |
| *Corymbia* | 5.6% | [3.3%, 8.7%] |
| *Darwinia* | 18.5% | [10.1%, 30.7%] |
| *Decaspermum* | 59.6% | [58.4%, 61.7%] |
| *Eucalyptus* | 21.9% | [18.3%, 27.2%] |
| *Eugenia* | 35.6% | [31.3%, 40.0%] |
| *Gossia* | 40.5% | [36.3%, 45.4%] |
| *Homoranthus* | 19.2% | [12.3%, 26.9%] |
| *Hypocalymma* | 4.8% | [0.2%, 13.1%] |
| *Lenwebbia* | 32.7% | [23.7%, 48.3%] |
| *Leptospermum* | 13.6% | [9.8%, 18.2%] |
| *Lindsayomyrtus* | 4.9% | [0.2%, 12.8%] |
| *Lithomyrtus* | 5.3% | [0.3%, 13.0%] |
| *Lophostemon* | 2.4% | [0.1%, 6.5%] |
| *Melaleuca* | 22.5% | [17.5%, 28.5%] |
| *Metrosideros* | 5.9% | [3.3%, 9.6%] |
| *Mitranthia* | 18.6% | [10.6%, 30.2%] |
| *Myrciaria* | 4.7% | [0.2%, 12.9%] |
| *Myrtus* | 24.0% | [11.0%, 51.1%] |
| *Neofabricia* | 10.1% | [0.4%, 29.0%] |
| *Osbornia* | 5.2% | [0.3%, 12.8%] |
| *Pilidiostigma* | 14.0% | [6.6%, 23.8%] |
| *Rhodamnia* | 36.8% | [32.1%, 42.5%] |
| *Rhodomyrtus* | 28.1% | [22.2%, 34.7%] |
| *Ristantia* | 15.2% | [6.7%, 29.0%] |
| *Sphaerantia* | 18.6% | [9.8%, 30.7%] |
| *Stockwellia* | 38.2% | [25.9%, 53.8%] |
| *Syzygium* | 13.0% | [10.0%, 16.1%] |
| *Thryptomene* | 10.9% | [0.4%, 29.3%] |
| *Tristania* | 18.8% | [9.8%, 30.7%] |
| *Tristaniopsis* | 21.8% | [14.4%, 31.1%] |
| *Uromyrtus* | 11.9% | [6.1%, 19.5%] |
| *Xanthostemon* | 27.0% | [19.0%, 35.6%] |

### 3.3. Myrtle rust infection risk under present and future climate

Under ‘present’ (2014) climate conditions based on the ACCESS-CM2 model and a conservative infection risk threshold (*I*_risk_ > 0.5) assuming lesser spread, suitable areas for myrtle rust are primarily concentrated along the eastern coastal and subcoastal regions, with smaller pockets in the southwest and southeast (Fig. 4). ‘Future’ (2100) climate projections show a southward shift in these suitable areas, particularly under more severe warming scenarios such as SSP3-7.0 (Fig. A.13).

**Figure 4.**
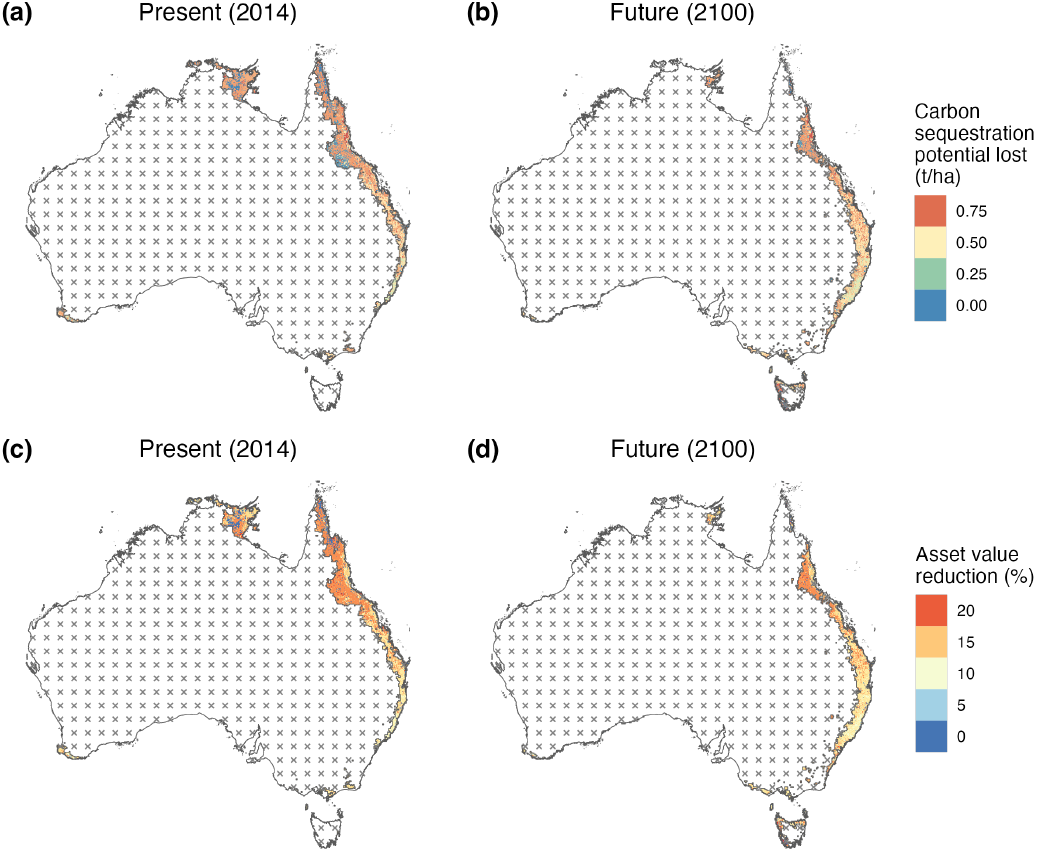
Projected impact of myrtle rust (exotic strains) on carbon sequestration and economic value. Estimated median annual reduction in carbon sequestration by woody plants due to myrtle rust under (a) present-day (2014) and (b) future (2100) climate conditions. Panels (c) and (d) show the corresponding annual percentage reduction in the economic value of sequestration for present and future conditions, respectively. Hashed areas indicate regions deemed climatically unsuitable for myrtle rust, based on our more conservative infection risk threshold (*I*_risk_ > 0.5, assuming less spread) derived from bioclimatic variables projected by the ACCESS-CM2 model, using the moderate emissions scenario SSP2-4.5 for future projections.

Similar patterns were observed across the two other CMIP6 climate models at the *I*_risk_ > 0.5 threshold (Fig. A.13). When applying a less conservative threshold (*I*_risk_ > 0.2, assuming greater spread), additional inland eastern areas and southwestern coastal/subcoastal regions were identified as suitable, both presently and under future conditions (Fig. A.12). However, these projections showed greater variability across models and scenarios. Detailed infection risk maps and bioclimatic drivers are provided in Appendix A.3.

### 3.4. Carbon sequestration reduction and dollar loss due to myrtle rust

Under present (2014) climate conditions (ACCESS-CM2, risk threshold > 0.5), myrtle rust (exotic strains) is estimated to reduce annual carbon sequestration by 1.6% (95% CI: 1.3%– 2.0%), with the greatest losses concentrated in northeastern Queensland (Fig. 4 a, c). This equates to an estimated annual economic loss of AUD $340 million [$284M, $422M].

Future projections (2100) under all SSP scenarios indicate slightly lower impacts. For instance, under the intermediate SSP2-4.5 scenario, carbon losses fall to 1.1% [0.9%, 1.3%], with annual economic losses of AUD $268 million [$224M, $322M]. This reduction is driven by the southward shift in suitable areas, which tend to have lower per-hectare carbon sequestration potential and economic value (Fig. 4 b, d).

Estimates for alternative climate models and the *I*_risk_ > 0.2 infection threshold (assuming greater spread of myrtle rust) are included in Appendix A.6. In general, results at the > 0.5 threshold were consistent across climate models. However, using the more inclusive > 0.2 threshold significantly increased and added more variability to projected losses (Fig. 5), with annual damages under future conditions (year 2100) reaching up to AUD $702 million [$588M, $872M] and a corresponding 5.5% [24.6%, 6.9%] reduction in national carbon sequestration under the ACCESS-CM2 climate model and intermediate emissions scenario (SSP2-4.5).

**Figure 5.**
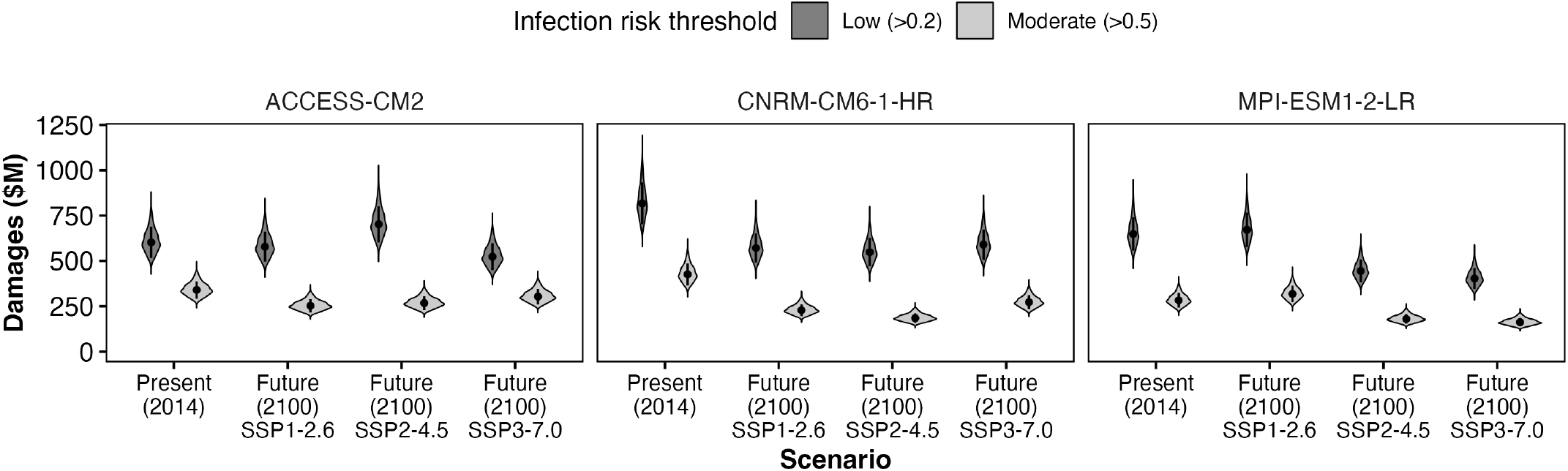
Estimated economic losses (i.e., ‘damages’ in 2015 AUD $M) from myrtle rust-induced reductions in carbon sequestration potential under ‘present’ (year 2014) and ‘future’ (year 2100) climate conditions. Losses are shown across two infection risk thresholds (*I*_risk_ > 0.2 and > 0.5, corresponding to assumed greater and lesser spread respectively), three climate models (ACCESS-CM2, CNRM-CM6-1-HR, MPI-ESM1-2-LR), and three future emissions scenarios (SSP1-2.6: low, SSP2-4.5: moderate, SSP3-7.0: high). Points indicate median estimates; error bars represent 95% interquartile ranges.

## 4. Discussion

Here, we used a data-driven modelling approach to estimate the possible nation-wide impact of myrtle rust on carbon sequestration in Australia and found that myrtle rust could have a potentially large effect on the environment — with an estimated dollar loss of over AUD $340 million on carbon sequestration assuming current climate conditions (estimated under the ACCESS-CM2 model), which could mean not being able to sequester the CO_2_ being emitted from over 5 Victorian coal power stations. We also found that under some of the more extreme future climate scenarios, carbon sequestration reductions could be less severe. This is because of reductions in future myrtle rust suitability in these scenarios caused by more extreme temperature and precipitation regimes — though, note that there were also other scenario forecasts with increased impacts of up to $702 million [$588M, $872M].

Our work shows that myrtle rust alone can have a sizeable impact on carbon sequestration, suggesting that the impact of invasive pests and diseases to ecosystem services may be systematically undervalued. Given that impacts to carbon sequestration are often tied with impacts to other ecosystem services (e.g. soil and air mediation, erosion control etc.), this provides additional evidence that invasive pests and diseases have big impacts to the environment. With the increased spread of invasive pests and diseases worldwide, systematic compilation of potential impacts and costs — to support biosecurity decision-making and climate-change related initiatives — is more critical than ever. Our method could be used to comprehensively and systematically estimate the impacts of invasive pests and diseases.

The main limitation of our modelling approach for comprehensive estimation of ecosystem impacts is its high data requirements. We had no plot data for some major vegetation groups in Australia, leading to underestimation of the potential impact of myrtle rust. Many MVGs were not sampled adequately and were represented by less than three plots, some of which were well below one hectare in size (e.g., MVG 1’s “Rainforests and Vine Thickets” total plot area was 0.08 ha), which could lead to either over- or under-estimation. While broad-scale remote sensing data exists, it is unable to resolve genus composition or capture genus-specific plant size distributions. These data gaps could be addressed by improving existing data public availability, conducting plot surveys in areas with high levels of uncertainty, and using remote sensing data to estimate biomass while making some simplified assumption about the overall impact due to invasive pests and diseases. Host distribution models (with plant size information) could also be incorporated, though this could affect the feasibility of using this workflow to comprehensively estimate impacts.

Our approach also has data gaps in regards to how myrtle rust impacts the carbon sequestration ability of individual plants. The majority of plant genera did not have quantitative information on the impact of myrtle rust, leading us to develop Bayesian estimates of impact based on qualitative susceptibility scores. In addition, the quantitative information that did exist considered different forms of impacts within different studies (e.g. branch dieback, leaf foliage number, height growth differences). Lastly, modelling carbon contribution of plants or impact on carbon sequestration can be done in a much more precise (and complex) manner than we have done. We focused on estimating the above-ground and below-ground biomass woody plants by genus, and did not include factors such as species, tree age, climate, or nutrient availability. While growth and impact could be modelled to account for these factors, data to support this (e.g., age and climate specific growth rates) was not available for many plant species. Hence, simplified assumptions on plant growth and pest/disease impact had to be made. Furthermore, there are other carbon contribution mechanisms involved (e.g., soil carbon sequestration by roots, carbon emissions from bushfire events) that can be considered in future studies that extend this work.

Note that our approach provides an estimate of long-term impacts and aims to examine the cost of failing to prevent an introduction rather than understand spatial dynamics of the invasion process. We approximate a “worst case” scenario of myrtle rust invasion in which it is instantaneously present across the areas in Australia which it is climatically suitable. Our approach does not model accrued impacts over time as the pathogen spreads and does not account for the likely scenario of a reactive management response that would slow or control the spread of the pathogen and reduce the impact of myrtle rust. It also does not consider host tree range shifts throughout the forecasting horizon [81]. Conversely, our approach focused on a decline in carbon capture ability, and therefore did not estimate the future emissions caused by the decomposition of trees killed by myrtle rust, which would further reduce net carbon sequestration and increases the effective potential damage caused by myrtle rust. Biosecurity measures and systems help protect the natural environment by reducing the risk of incursion of invasive pests and diseases, providing surveillance to detect incursions, and post-border management to deal with incursions. Given the impact of invasive pests and diseases on the carbon sequestration, our work shows the importance of biosecurity to delivering on climate-change related commitments, including net-zero emissions targets and reforestation efforts, while also providing sufficient granularity to be used as an input to support biosecurity decision-making. We are not aware of any other work that links invasive pathogens quantitatively to climate mitigation impacts — and expect many high profile forest pests and pathogens to cause similar damages (e.g., Emerald Ash Borer, *Agrilus planipennis*, [82]). Given the present undervaluation of biodiversity and potential environmental consequences given invasive pests and diseases, relative to impacts to agriculture and other primary industries, there needs to be further work in systematically (re)estimating ecosystem impacts to truly understand the magnitude of consequences that we could be facing. We provide the methodology herein to estimate these ecosystem system impacts and to incorporate traditionally non-market impacts of invasive species into economic impact assessments.

## 5. Conclusion

Increasing native ecosystems’ capacities to act as carbon sinks is a major tool for climate change mitigation, and trees perform important climate change adaptation services, especially in cities under a warming climate [83]. However, planted and existing trees are susceptible to a range of pests and diseases. Our modelling workflow employs existing information about invasive species impact at the host level to estimate the impact at the country-wide level. We found that myrtle rust could have a sizeable impact on carbon sequestration in Australia, in particular due to their impacts on *Eucalyptus* tree, which play a major role in sequestering carbon and are also very susceptible to myrtle rust. While this case study has focussed on myrtle rust and carbon sequestration, the methodology is transferable across invasive species and ecosystem services, allowing for systematic estimation of the potential impacts and costs which can then support decision-making in biosecurity and climate-change initiatives.

## CRediT authorship contribution statement

**T.P. Le:** Conceptualization, Data curation, Formal Analysis, Funding acquisition, Investigation, Methodology, Writing – original draft, Writing – review & editing. **M. Theng:** Data curation, Formal Analysis, Investigation, Methodology, Visualization, Writing – review & editing. **C.M. Baker:** Funding acquisition, Project administration, Writing – review & editing. **I.R. Abell:** Writing – review & editing. **T. Kompas:** Funding acquisition, Writing – review & editing. **E. Hudgins:** Funding acquisition, Methodology, Writing – review & editing, Investigation.

## Declaration of competing interest

The authors declare that they have no known competing financial interests or personal relationships that could have appeared to influence the work reported in this paper.

### Acknowledgements

We acknowledge the input and advice provided Pip Griffin and Mirja Guldner.

## Data availability

Carbon sequestration value per ha per MVG across Australia were obtained from [7]. Datasets with species and diameter information of each plant record, and coordinates of the plot (or site) in Australia can be found: Biomass Plot Library [45], Calperum Mallee Stem Diameter, Height and Aboveground Woody Biomass Data [62], Robson Creek Rainforest Diameter, Height and Aboveground Woody Biomass Data [63]. Myrtle rust impacts to individual species with references are collated in Table A.3.

## Appendix A. Supplementary

### Appendix A.1. Spatial distribution of trees and shrubs - data sources

We collated vegetation plot datasets of individual trees and shrubs sampled from a range of vegetation types. These datasets contained species and diameter information of each plant record, and coordinates of the plot (or site):

- Biomass Plot Library [45], which contains a national collation of stem inventory data and biomass estimation of 1,467 tree and shrub species across 12,663 field sites across most bioregions in Australia;
- Calperum Mallee Stem Diameter, Height and Aboveground Woody Biomass Data [62], which contains stem diameter, height measurement and above ground living biomass calculations for a Mallee Eucalypt dominated woodland (2015 – present);
- Robson Creek Rainforest Diameter, Height and Aboveground Woody Biomass Data [63], which contains stem diameter, height measurement and above ground living biomass calculations for an Australian tropical rainforest derived from a comprehensive ground survey of a large area (2009 – present);

Note that we discarded records before 2013 and assumed that the dataset was representative of the present vegetation within the surveyed plots. In addition, we discarded records without species or diameter information, and only kept the latest record for individuals that were recorded more than once. This resulted in an inventory of 28,355 unique trees across 159 survey plots, covering 50.5 hectares and representing 13 of the 32 Major Vegetation Groups (Table A.2).

#### Appendix A.2. Biomass estimation

Both above-ground biomass (AGB) and below-ground biomass (BGB) contribute to the woody vegetation carbon sink. Allometric models provide cost-effective methods for quantifying sequestration of biomass carbon from woody vegetation. Ground-based estimates are typically obtained by applying these models to field measurements of biometric data such as stem diameter or plant height. The following subsections detail the functional grouping and models specific to AGB and BGB estimation.

##### Appendix A.2.1. Above-ground biomass (AGB)

To estimate AGB, individual tree and shrub records were first grouped into the following plant functional types [64]:

- *F*_Shrub_: shrubs or small trees characterized by being relatively short (generally < 2 m height) and typically multistemmed or highly branched, with a relatively small (< 7 cm) stem diameter;
- *F*_Multi_: multistemmed hardwood (angiosperm) trees, including mallees from the genus *Eucalyptus*, and trees from the genus *Acacia*;
- *F*_Euc_: typically single-stemmed hardwood trees from the genus *Eucalyptus* and closely related genera of *Corymbia* and *Angophora*;
- *F*_Other-H_: other tree species that typically have single stems and relatively high wood density (mean 0.67 g cm^™3^);
- *F*_Other-L_: other trees, namely conifers from the genera of *Pinus, Araucaria* and *Agathis*, that typically have single stems and relatively low stem wood density (mean 0.40 g cm^™3^).

The respective models developed for AGB prediction (kg) were then applied to each tree or shrub [64]:

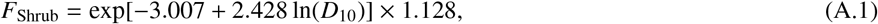

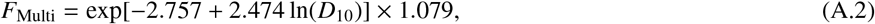

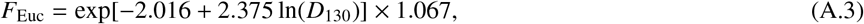

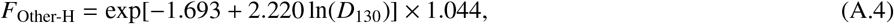

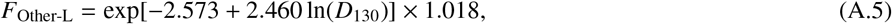

where *D*_10_ is the tree diameter measured 10 cm from the ground and *D*_130_ is tree diameter measured 130 cm from the ground.

##### Appendix A.2.2. Below-ground biomass (BGB)

To estimate BGB, individual tree and shrub records were first grouped into the following plant functional types [65]:

- *F*_Shrub,Ac_: shrubs and small multi-stemmed trees.
- *F*_Mallee_: multi-stemmed (mallee) trees from the genus *Eucalyptus*, which commonly have a lignotuber and relatively high wood density (mean density of 0.88 g cm^™3^).
- *F*_Tree_: typically single-stemmed trees of relatively high wood density (mean density of 0.69 g cm^™3^).
- *F*_Radiata_: the specific tree species *Pinus radiata*, of relatively low stem woody density (mean density of 0.40 g cm^™3^). The respective models developed for BGB prediction (kg) were then applied to each tree or shrub [65]:

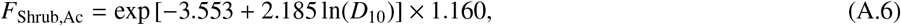

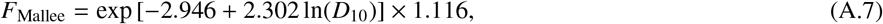

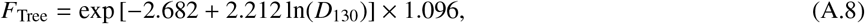

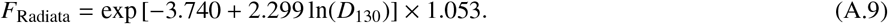

Each tree or shrub record (i.e., individual) *i* was assigned to an AGB and BGB functional group, and the respective allometric equations were applied to the diameter measurement to obtain the total biomass estimate (*B*_*i*_) for the record *i*:

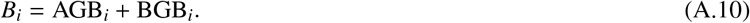

##### Appendix A.3. Myrtle rust infection risk

To calculate infection risk as per Beresford et al [72], we first define the response to infection risk, *Y*,

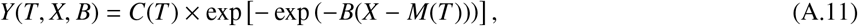

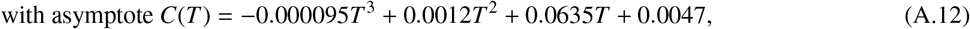

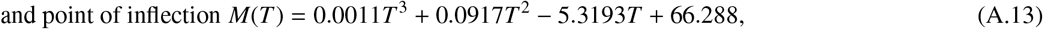

where we evaluate the infection risk *Y* on: 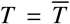, the mean temperature during high relative-humidity hours (relative humidity above 85%); *X*, the number of hours of high relative humidity (relative humidity above 85%) (Fig. A.9), and *B* = 0.39 is a fixed rate.

As we do not have hourly information regarding temperature and relative humidity, we estimate them based on known functional forms [73].

We estimate the hourly temperatures using a cosine function (Eq. (7) of [73]), assuming maximum temperatures occur at 3 pm (hour *h*_*max*_ = 15) and minimum temperatures occur at 6 am (hour *h*_*min*_ = 6):

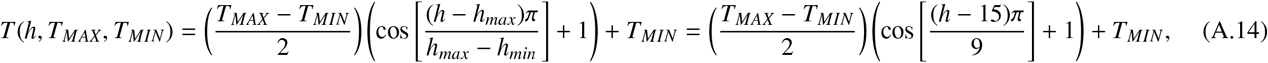

where *h* is the hour at which we are calculating the temperature and we take *T*_*MAX*_ to be the mean of the maximum recorded temperatures across a year (Fig. A.6) and *T*_*MIN*_ to be the mean of the minimum recorded temperatures (Fig. A.7) across a year.

We estimate hourly relative humidity *RH*(*h, d*), on given day *d*, based on the daily actual vapour pressure *V*(*d*) and the hourly saturation vapour pressure *P*(*h, d*):

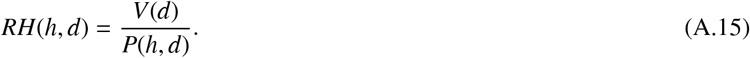

The hourly saturation vapour pressure is estimated using the Tetens equation:

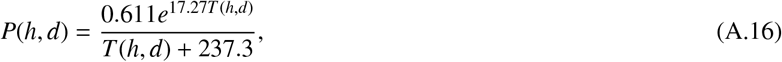

where *T* (*h, d*) is the temperature from Eq. (A.14). The daily actual vapour pressure *V*(*d*) is estimated from the daily humidity data and the mean saturation vapour pressure on that day *d*, 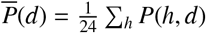:

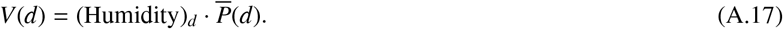

This allows us to calculate *X*, the number of high RH hours in each day based on when *RH*(*h, d*) > 0.85.

Next, we define the light intensity *L*, which is dependant on the hourly solar radiation *S R* data (Fig. A.8):

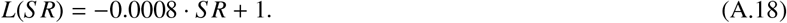

Finally, infection risk is calculated as:

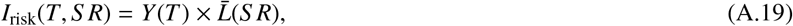

where 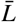 (*S R*) is the daily average of the hourly light intensity values *L*(*S R*).

Note that we calculate infection risk at both *T* = *T*_*MAX*_ (Fig. A.11) and *T* = *T*_*MIN*_ (Fig. A.10), as a conservative estimate to myrtle rust suitability. Raster pixels exceeding the threshold (evaluated at 0.2 and 0.5) for either *I*_risk_(*T*_*MAX*_, *S R*) or *I*_risk_(*T*_*MIN*_, *S R*) were classified as areas climatically suitable for myrtle rust. Fig. A.12 and Fig. A.13 show myrtle rust suitability areas when *I*_risk_ > 0.2 and *I*_risk_ > 0.5 respectively for multiple climate models.

**Figure A.6:**
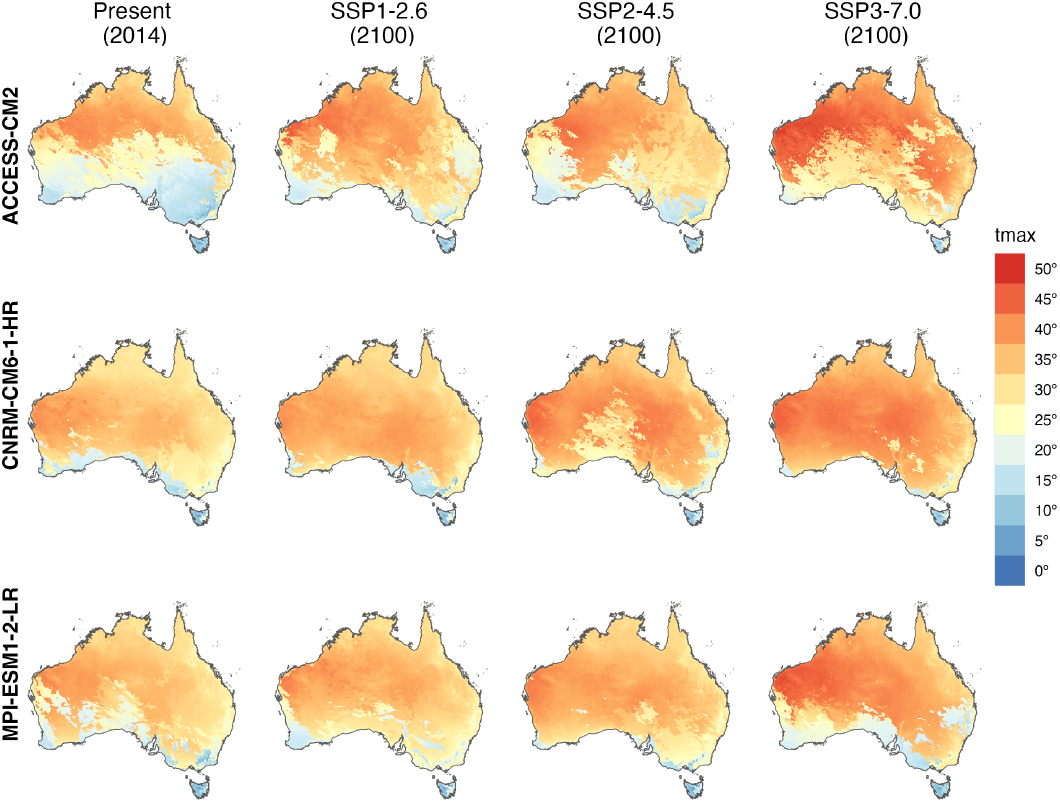
Maximum temperature of the wettest quarter given different CMIP6 climate models (ACCESS-CM2, CNRM-CM6-1-HR, MPI-ESM1-2-LR) under present conditions (year 2014) and three future (year 2100) climate scenarios (SSP1-2.6: low emissions, SSP2-4.5: moderate emissions, SSP3-7.0: high emissions).

**Figure A.7:**
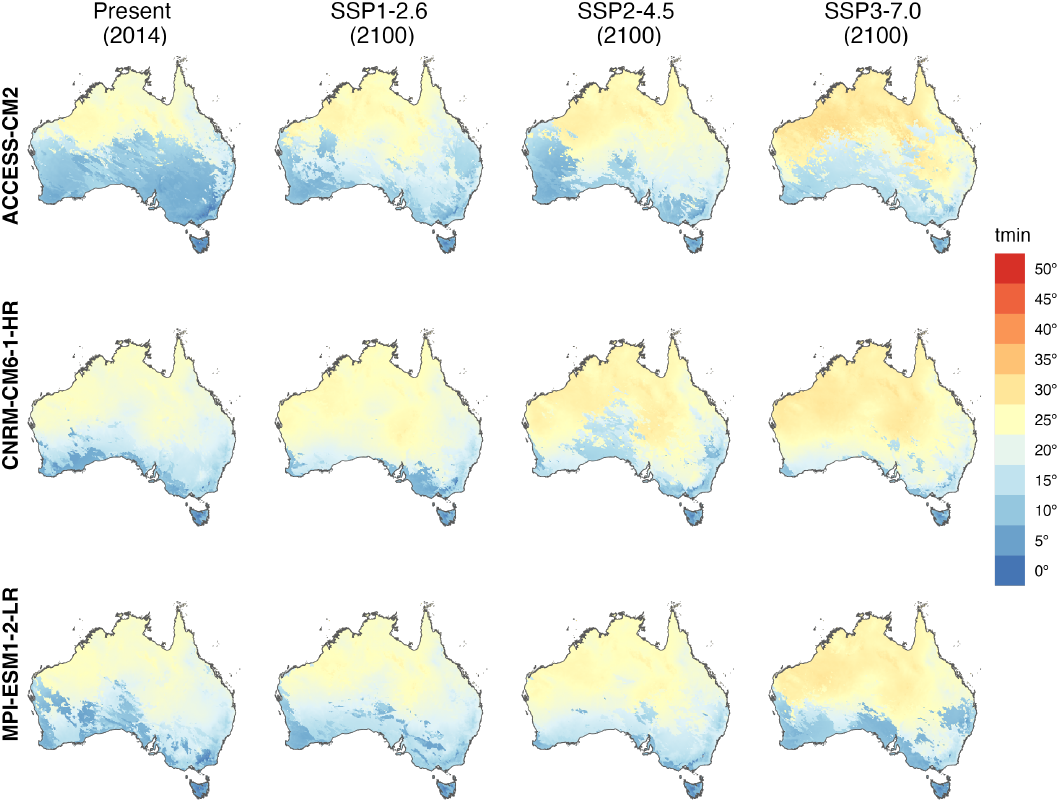
Minimum temperature of the wettest quarter given different CMIP6 climate models (ACCESS-CM2, CNRM-CM6-1-HR, MPI-ESM1-2-LR) under present conditions (year 2014) and three future (year 2100) climate scenarios (SSP1-2.6: low emissions, SSP2-4.5: moderate emissions, SSP3-7.0: high emissions).

**Figure A.8:**
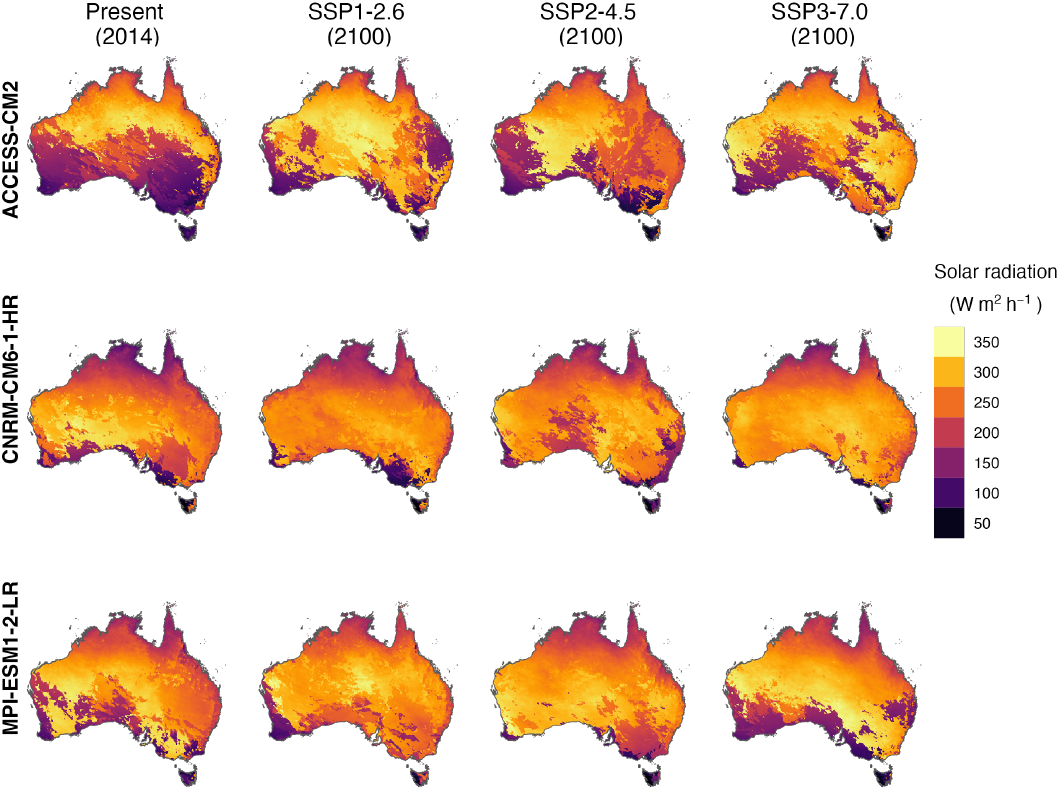
Solar radiation of the wettest quarter given different CMIP6 climate models (ACCESS-CM2, CNRM-CM6-1-HR, MPI-ESM1-2-LR) under present conditions (year 2014) and three future (year 2100) climate scenarios (SSP1-2.6: low emissions, SSP2-4.5: moderate emissions, SSP3-7.0: high emissions).

**Figure A.9:**
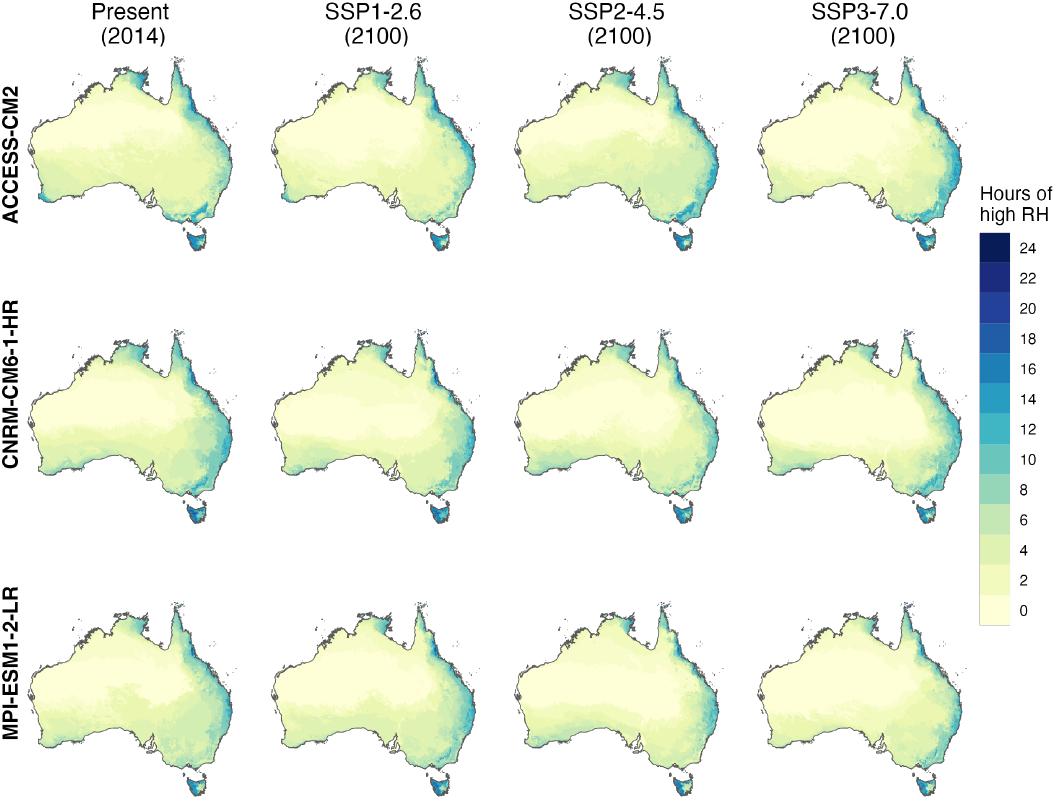
Hours of high relative humidity of the wettest quarter given different CMIP6 climate models (ACCESS-CM2, CNRM-CM6-1-HR, MPI-ESM1-2-LR) under present conditions (year 2014) and three future (year 2100) climate scenarios (SSP1-2.6: low emissions, SSP2-4.5: moderate emissions, SSP3-7.0: high emissions).

**Figure A.10:**
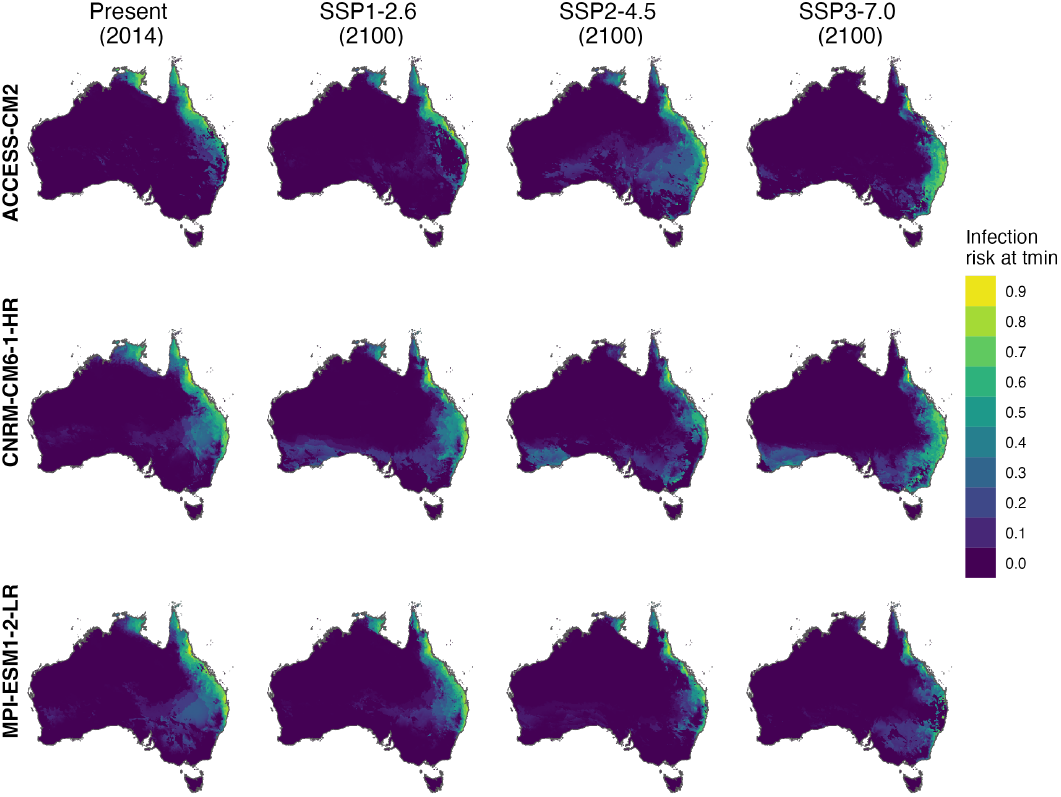
Projected myrtle rust infection risk at the minimum temperature of the wettest quarter, derived from bioclimatic variables under different CMIP6 climate models (ACCESS-CM2, CNRM-CM6-1-HR, MPI-ESM1-2-LR) under present conditions (year 2014) and three future (year 2100) climate scenarios (SSP1-2.6: low emissions, SSP2-4.5: moderate emissions, SSP3-7.0: high emissions).

**Figure A.11:**
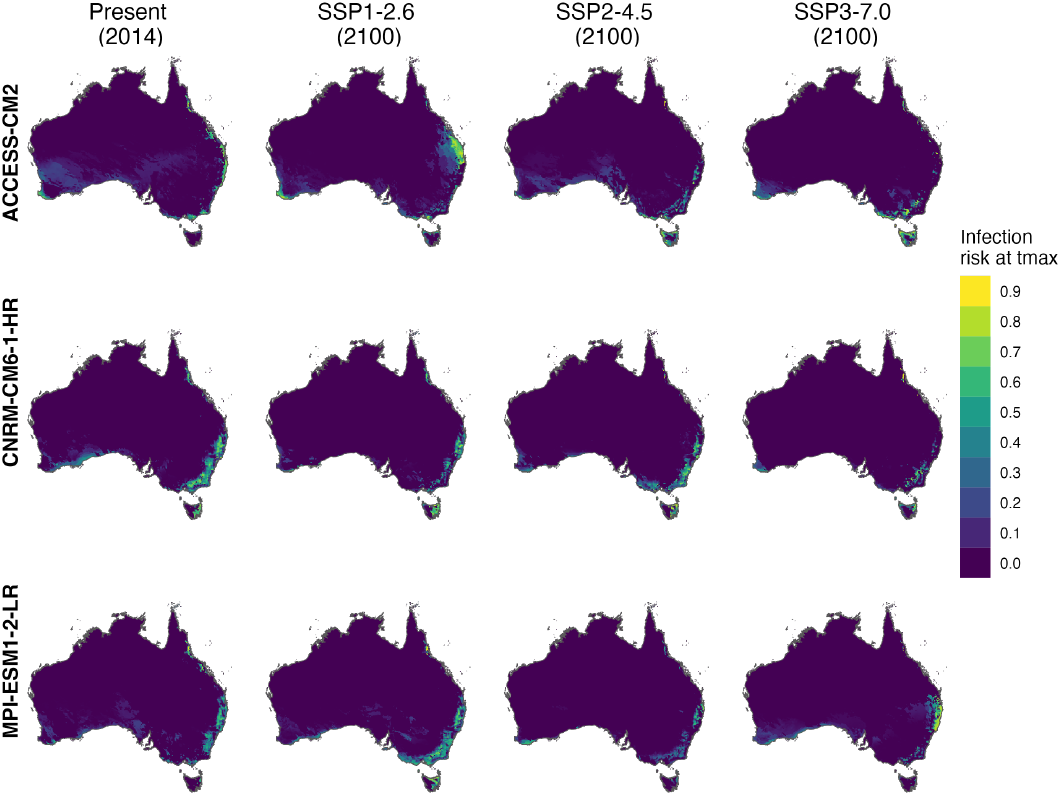
Projected myrtle rust infection risk at the maximum temperature of the wettest quarter, derived from bioclimatic variables under different CMIP6 climate models (ACCESS-CM2, CNRM-CM6-1-HR, MPI-ESM1-2-LR) under present conditions (year 2014) and three future (year 2100) climate scenarios (SSP1-2.6: low emissions, SSP2-4.5: moderate emissions, SSP3-7.0: high emissions).

**Figure A.12:**
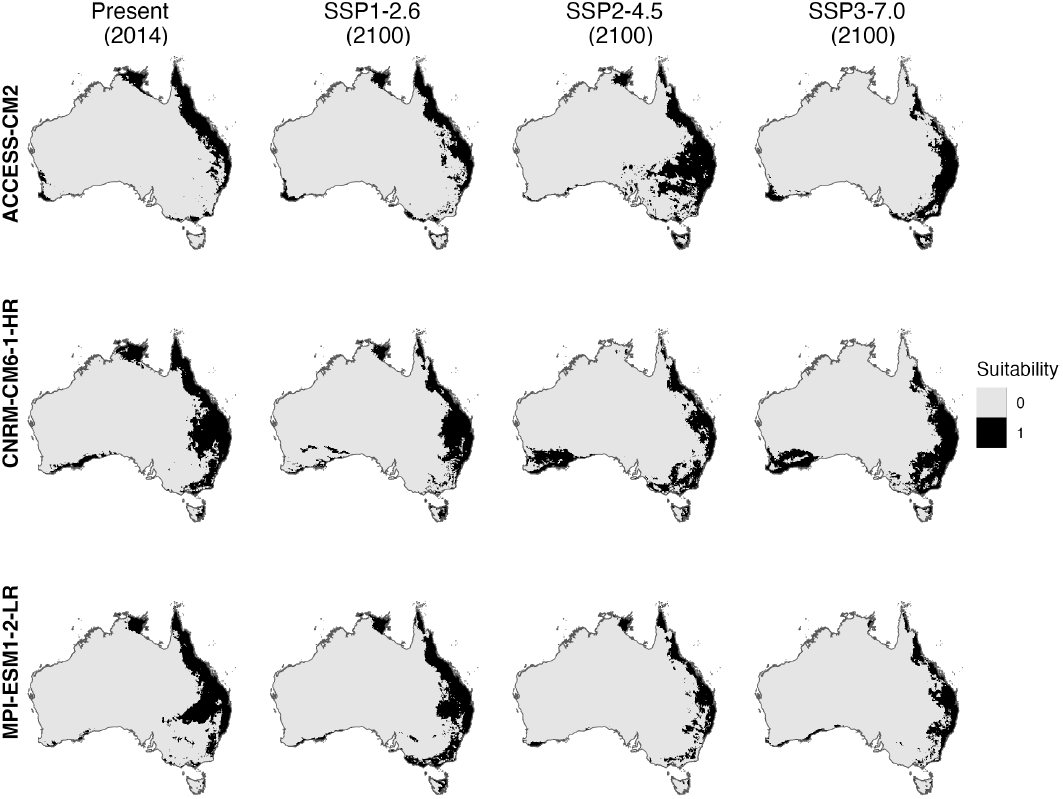
Binary map of myrtle rust infection risk (threshold > 0.2, assuming greater spread), derived by combining projected risk at minimum and maximum temperatures (see Fig. A.10 and Fig. A.11). Results are shown for three CMIP6 climate models (ACCESS-CM2, CNRM-CM6-1-HR, MPI-ESM1-2-LR) under present conditions (year 2014) and three future (year 2100) climate scenarios (SSP1-2.6: low emissions, SSP2-4.5: moderate emissions, SSP3-7.0: high emissions). We assume that myrtle rust (exotic strains) can only spread to areas in black (*suitability* = 1).

**Figure A.13:**
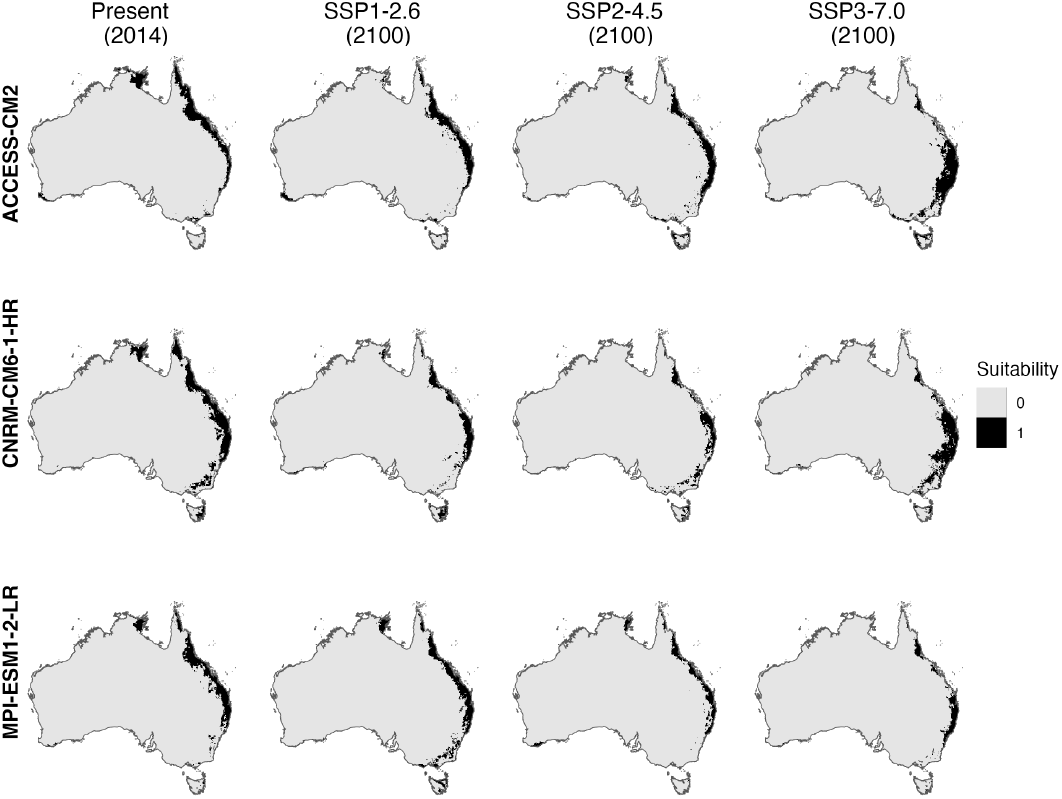
Binary map of myrtle rust infection risk (threshold > 0.5, assuming lesser spread), derived by combining projected risk at minimum and maximum temperatures (see Fig. A.10 and Fig. A.11). Results are shown for three CMIP6 climate models (ACCESS-CM2, CNRM-CM6-1-HR, MPI-ESM1-2-LR) under present conditions (year 2014) and three future (year 2100) climate scenarios (SSP1-2.6: low emissions, SSP2-4.5: moderate emissions, SSP3-7.0: high emissions). We assume that myrtle rust (exotic strains) can only spread to areas in black (*suitability* = 1).

#### Appendix A.4. Known myrtle rust impacts

Table A.3 lists the species with some known quantitative information regarding myrtle rust impacts, and their source.

**Table A.3:** Species with known quantitative myrtle rust impacts.

| Species | Damage | Information |
| --- | --- | --- |
| *Acmena smithii* (aka *Syzygium smithii*) | 0.1252 | Branch dieback, averaged across size classes [84] |
| *Angophora costata* | 0.1176 | Height difference between control and inoculated plants [53] |
| *Archirhodomyrtus beckleri* | 0.9428 | Branch dieback, averaged across size classes [84] |
| *Baeckea linifolia* | 0.11 | Reduction in leaf number [85] |
| *Callistemon* (= *Melaleuca*) *citrinu* | 0.2419 | Height difference between control and inoculated plants [53] |
| *Callistemon* (= *Melaleuca*) *rigidus* | 0.2593 | Height difference between control and inoculated plants [53] |
| *Corymbia citriodora* | 0.1212 | Growth rate reduction between control and inoculated plants [86] |
| *Corymbia maculata* | 0 | Not susceptible [87] |
| *Decaspermum humile* | 0.7778 | Branch dieback, averaged across size classes [84] |
| *Eugenia reinwardtiana* | 0.5 | Defoliation [26] |
| *Eugenia koolauensis* | 0.7 | Mortality of species in Hawaii [26] |
| *Eucalyptus grandis* × *Eucalyptus urophylla* clone | 0.89 at 20 months,<br>0.34 at 5 years | Growth difference with and without a preventive rust treatment [88] |
| *Eucalyptus grandis* | 0.23 | Wood volume losses in plantation [89] |
| *Eucalyptus globulus* | 0 | Not susceptible [87] |
| *Eucalyptus globoides* | 0.0962 | Height difference between control and inoculated plants [53] |
| *Eucalyptus moluccana* | 0.2097 | Height difference between control and inoculated plants [53] |
| *Eucalyptus viminalis* | 0 | No height difference between control and inoculated plants [53] |
| *Gossia hillii* | 0.9896 | Branch dieback, averaged across size classes [84] |
| *Leptospermum laevigatum* | 0.24 | Height reduction in seedlings [85] |
| *Lophostemon confertus* | 0 | Not susceptible [90] |
| *Melaleuca quinquenervia* | 0.45 | Reduction in biomass when infected [85] |
| *Melaleuca squarrosa* | 0 | Not susceptible [90] |
| *Melaleuca styphelioides* | 0.2315 | Height difference between control and inoculated plants [53] |
| *Metrosideros polymorpha* | 0.08 | Mortality rate in seedlings and young saplings [91] |
| *Rhodamnia maideniana* | 0.9671 | Branch dieback, averaged across size classes [84] |
| *Rhodamnia rubescens* | 0.12 | Mean tree death [52] |
| *Rhodomyrtus psidioides* | 0.57 | Mean tree death [52] |

#### Appendix A.5. Estimating myrtle rust impacts via Bayesian inference

Many plant genera had no quantitative information regarding the impact of myrtle rust. However, Makinson [26] had reviewed the literature and produced a summary table regarding the susceptibility of many species to myrtle rust infection. Figure A.14 shows a plot of the distribution of ratings from Makinson [26] — “relatively tolerant”, “moderate susceptibility”, “high susceptibility”, and “extreme susceptibility”.

In order to convert these ratings into numerical damages, we use Bayesian inference which can combine the quantitative damage information and qualitative susceptibility scores to estimate the potential impact of myrtle rust on species. In particular, we use Bayesian inference to estimate the numerical boundaries of each of the scores “relatively tolerant”, “moderate susceptibility” and so forth.

We first assume that the underlying probability density for damages occurred by myrtle rust across all plant species is a normal distribution that is truncated between [0,1] (corresponding to 0% damage to 100% damage)

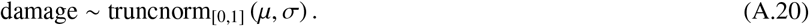

We will estimate the mean µ and standard deviation σ that characterises this distribution. The mean µ has a somewhat informative prior, where we assume that most plants are not very susceptible to myrtle rust, thus setting the midpoint at 0 (albeit with a broad standard deviation 2):

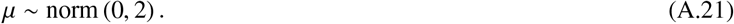

We assume a fairly broad uniform distribution on the standard deviation σ from 0 to 10, but not too broad as we do not expect a resultant uniform distribution for the damage density probability across all species (i.e. see Figure A.14):

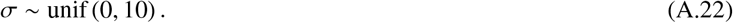

Next, we define the thresholds for the different susceptibility categories:

**Figure A.14:**
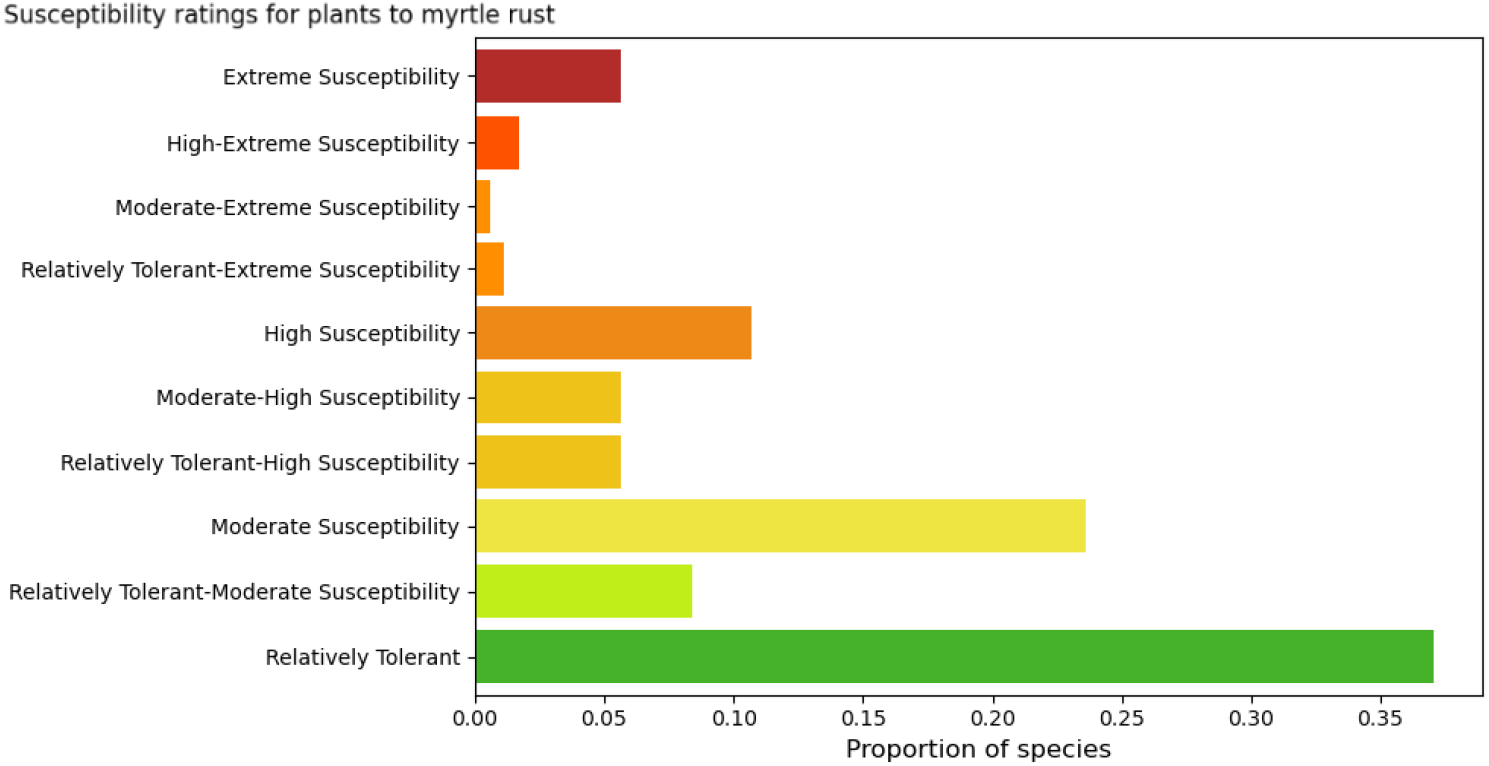
Distribution of plant susceptibility ratings to myrtle rust, from Makinson [26]. Note that some plant species were estimated to have a wide susceptibility range e.g, between Moderate-Extreme Susceptibility – these ranges were included during the Bayesian estimation of impact

- *θ*_RT_ refers to the relatively-tolerant (RT) threshold, where species with RT rating have a damage value between [0, *θ*_RT_];
- *θ*_MS_ refers to the moderate-susceptibility (MS) threshold, where species with a MS rating have a damage value between [*θ*_RT_, *θ*_MS_];
- *θ*_HS_ refers to the high-susceptibility (HS) threshold, where species with a HS rating have a damage value between [*θ*_MS_, *θ*_HS_];
- and finally, species with a extreme-susceptibility (ES) rating have a damage value between [*θ*_HS_, 1].

See Figure 2 in the main text for an example of how these thresholds play out.

We assign truncated normal priors on the thresholds. These are semi-informative, with the means located such that the interval [0,1] is evenly separated into four (corresponding the four different categories), with a moderate standard deviation:

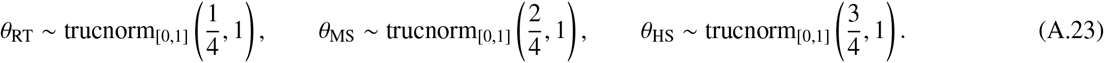

As we have two different data sources (quantitative data and qualitative scores) and no clear likelihood, we use Approximate Bayesian Computation with rejection sampling. This involves generating a set of parameter choices drawn from their prior distributions, simulating hypothetical datasets based on the drawn parameters, and comparing the simulated datasets with the data sources. If the generated datasets are sufficiently similar to the data sources, then we accept the parameter set. We repeat until we produce a posterior distribution of the parameters. In practice, we:

- We sample the candidate parameters 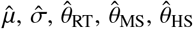 from their prior distributions. Note that during sampling, we also impose that *θ*_RT_ < *θ*_MS_ < *θ*_HS_.
- We consider genera with both known impact from myrtle rust *d*_*i*_ and susceptibility rating *r*_*i*_. We use the damage *d*_*i*_ and candidate thresholds 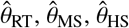 to predict their susceptibility rating 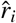:

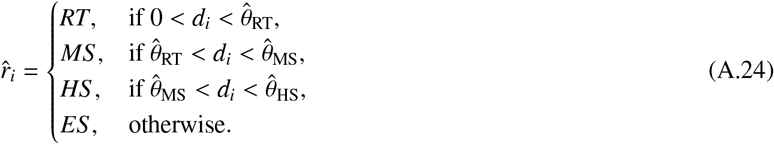

We then take the distance between the predicted susceptibility rating and the actual rating:

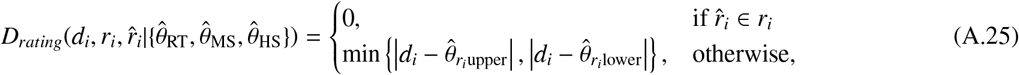

where 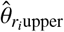 and 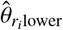 define the right and left thresholds for the correct rating. That is, the distance is zero if the predicted rating is correct, otherwise it is the distance from the damage *d*_*i*_ to its correct category.

We do this for all genera with combined known impact from myrtle rust and susceptibility rating and sum them:

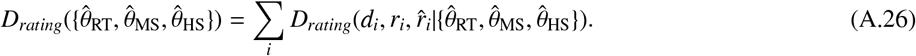

- Next, we consider the susceptibility rating information listed in Makinson [26]. Using the sampled parameters 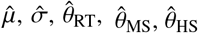, we estimate the distribution of outcome ratings across the different categories, 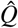. Then, we compare the simulated ratings with the actual ratings frequency *P*_data_ (i.e., of Fig. A.14). We use the Kullback–Leibler (KL) divergence to calculate the difference between the two:

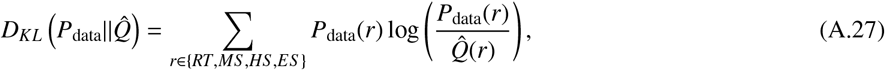

where *P*_data_(*r*) is the frequency of genera with rating *r* from the data, and 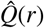 is the frequency of genera with the rating *r* from the generated data.

We use two thresholds for the two distance measures, accepting sampled parameters 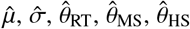if *D*_*rating*_ < 0.5 and *D*_*KL*_ < 0.1. Note that this leads to an acceptance rate of less than 0.1%. Two distinct acceptable thresholds are used for *D*_*rating*_ and *D*_*KL*_ due to their complex relationship: as shown in Figure A.15, *D*_*KL*_ varies across magnitudes from less than 10^™3^ to over 10^2^, while *D*_*rating*_ is mostly restricted between ≈ 0.4 and 2.7. Using a combined distance of *D*_*rating*_ · *D*_*KL*_ will overwhelmingly favour small values of *D*_*KL*_ whilst ignoring the outcome of *D*_*rating*_.

We repeat this process until we have accepted 20000 parameter sets, which form a joint posterior distribution of the parameters. The marginal posterior distributions of the parameters are shown in Figures A.16 and A.17. We then use the posterior parameter distribution to create 1000 unique estimations for myrtle rust damage values for various genera, which is presented as median and 95% interval in Table 1. Genera with no information were assumed to not be susceptible to myrtle rust, as a conservative estimate.

**Figure A.15:**
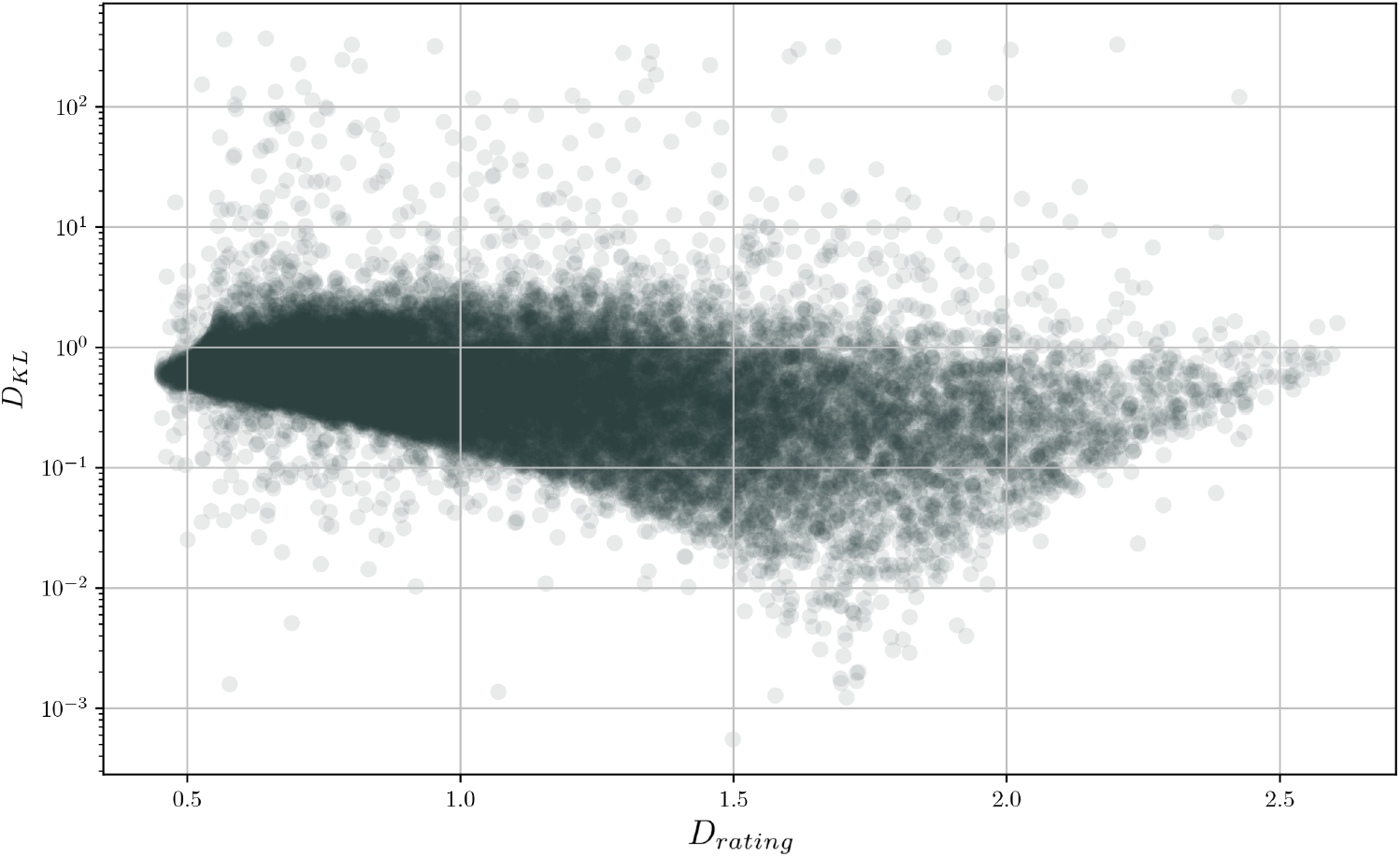
Plot of the relationship between Eq. (A.26) and Eq. (A.27)

**Figure A.16:**
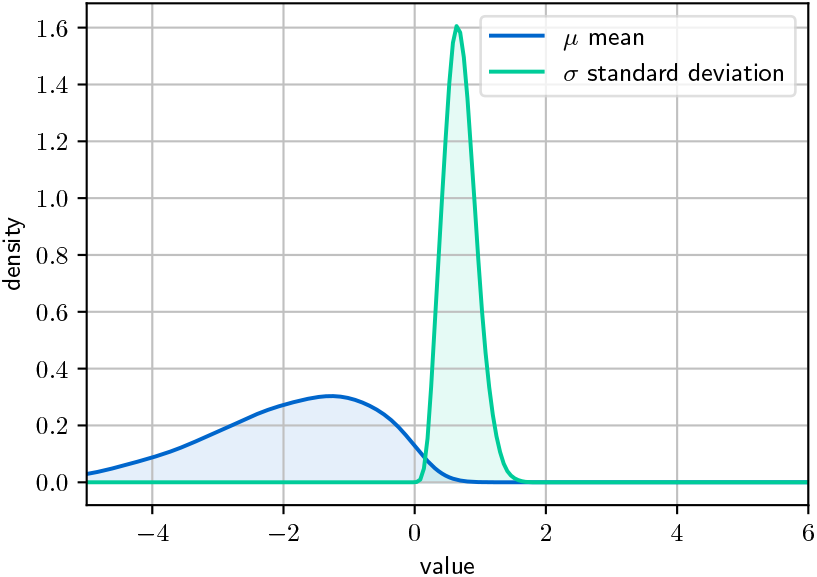
The posteriors for the parameters µ and σ, which define the truncated normal shape of the myrtle rust damages distribution (Eq. (A.20)).

**Figure A.17:**
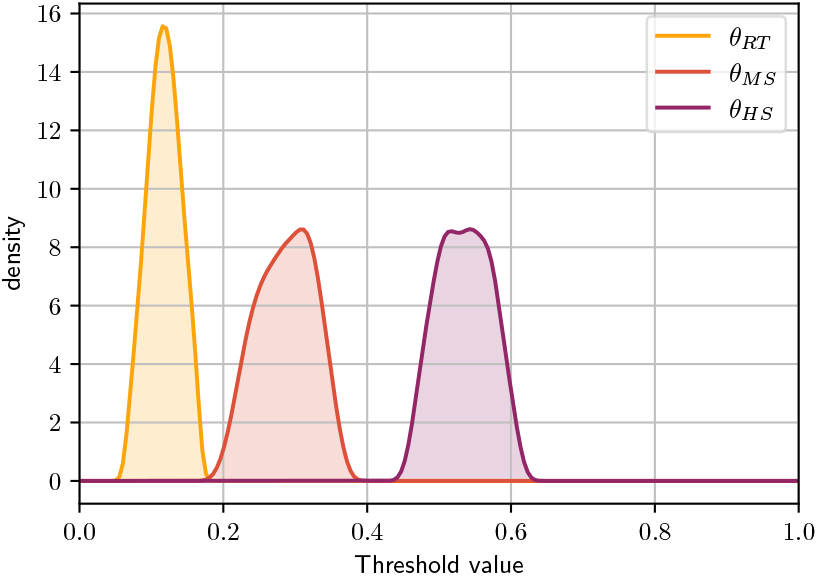
The posteriors of the parameters θ_RT_, θ_MS_, and θ_HS_, which define the thresholds for the upper boundary of myrtle rust damages to relatively tolerant, moderately susceptible, and highly susceptible genera, respectively.

#### Appendix A.6. Extended results

Summary estimates of annual reductions in carbon sequestration potential and associated dollar losses due to myrtle rust, under each climate model, emissions scenario (for future projections), and infection risk threshold, are presented in Table A.4.

**Table S4:**
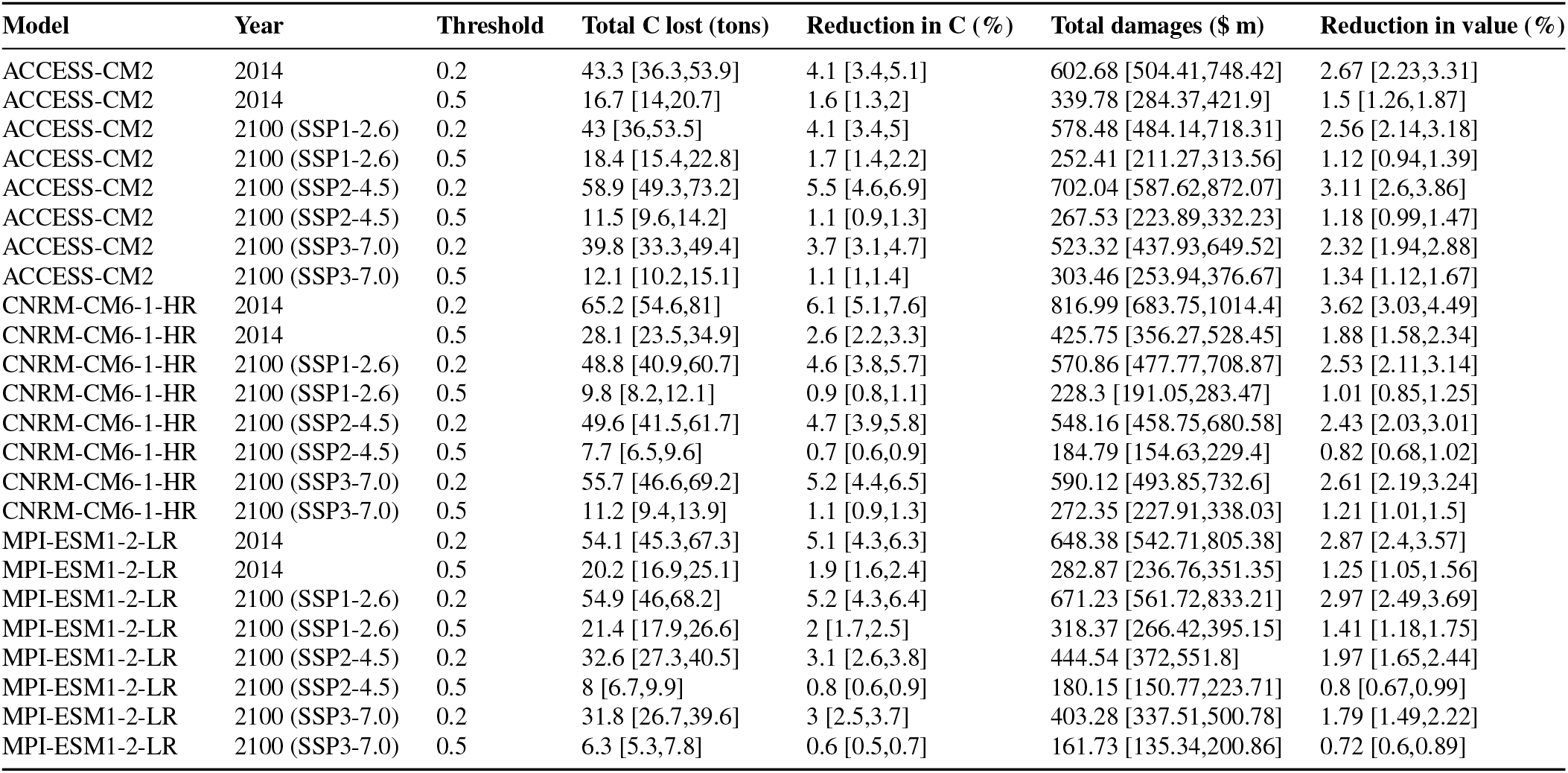
Summary estimates of the annual impact of myrtle rust on carbon sequestration potential, including total carbon lost (‘Total C lost (tons)’), percentage reduction in carbon sequestration (‘Reduction in C (%)’), and associated economic damages in 2015 AUD millions (‘Total damages ($m)’). Results are disaggregated by CMIP6 climate model (‘Model’), projection year (‘Year’), infection risk threshold (‘Threshold’), and emissions scenario for the year 2100 (SSP1-2.6: low emissions; SSP2-4.5: moderate emissions; SSP3-7.0: high emissions).

## Footnotes

1 Note, the database was queried first by filtering the ‘Environment’ field containing ‘Terrestrial’ or ‘Diverse/Unspecified’ and ‘Impacted sector’ field containing ‘Environment’. Under this filtering, there were 146 entries, of which 97 we could_source (the others were not accessible or not in English). Of these 97, 19 mentioned ecosystem services or carbon sequestration. Note that invasive pest impacts to ecosystem services are not all filed under ‘Impacted sector=Environment’ in the InvaCost database.

